# Direct Antimicrobial Resistance Prediction from clinical MALDI-TOF mass spectra using Machine Learning

**DOI:** 10.1101/2020.07.30.228411

**Authors:** Caroline Weis, Aline Cuénod, Bastian Rieck, Felipe Llinares-López, Olivier Dubuis, Susanne Graf, Claudia Lang, Michael Oberle, Maximilian Brackmann, Kirstine K. Søgaard, Michael Osthoff, Karsten Borgwardt, Adrian Egli

## Abstract

Early administration of effective antimicrobial treatments is critical for the outcome of infections. Antimicrobial resistance testing enables the selection of optimal antibiotic treatments, but current culture-based techniques take up to 72 hours. We have developed a novel machine learning approach to predict antimicrobial resistance directly from MALDI-TOF mass spectra profiles of clinical samples. We trained calibrated classifiers on a newly-created publicly available database of mass spectra profiles from clinically most relevant isolates with linked antimicrobial susceptibility phenotypes. The dataset combines more than 300,000 mass spectra with more than 750,000 antimicrobial resistance phenotypes from four medical institutions. Validation against a panel of clinically important pathogens, including *Staphylococcus aureus*, *Escherichia coli*, and *Klebsiella pneumoniae*, resulting in AUROC values of 0.8, 0.74, and 0.74 respectively, demonstrated the potential of using machine learning to substantially accelerate antimicrobial resistance determination and change of clinical management. Furthermore, a retrospective clinical case study found that implementation of this approach would have resulted in a beneficial change in the clinical treatment in 88% (8/9) of cases. MALDI-TOF mass spectra based machine learning may thus be an important new tool for antibiotic stewardship.

---

Antimicrobial resistant bacteria and fungi pose a serious and increasing threat to the achievements of modern medicine^1, 2^. Infections with antimicrobial resistant pathogens are associated with substantial morbidity, mortality, and healthcare costs^3^. Rapid treatment with an effective antimicrobial is critical for the outcome of an infection^4, 5^. However, antimicrobial therapy and dosage need to account for the resistance profiles of presumed pathogens, and also have to consider host-specific factors such as patient age, kidney function, and previous medical history. Early identification of the microbial species causing an infection can allow potential therapeutic options to become more targeted based on e.g. intrinsic resistance mechanisms and local epidemiology of resistance^6, 7^. However, only a detailed resistance profile permits treatments to be fully optimised. With current culture-based methods, the time from sample collection to resistance reporting can take up to 72 hours, meaning that for a substantial period, a patient may be receiving a too narrow- or too broad-spectrum antimicrobial drug^8, 9^. To limit the infection-related risk to a patient, broad-spectrum antibiotics are very often used. The concept of an optimal selection of an antibiotic drug is an important pillar of antibiotic stewardship and has gained significant attention owing to the global emergence and spread of antibiotic resistant pathogens. A reduction in the time required for a resistance profile to become available will not only substantially improve patient outcomes, but would also align well with other goals of antibiotic stewardship^10^, including reducing reliance on precious broad-spectrum antibiotic treatments, reducing unnecessary broad antibiotic use, and thereby combating the development of antibiotic resistance. In addition, rapid information on antimicrobial resistance may help to speed up infection prevention measures such as the isolation or cohorting of patients infected with presumed multidrug resistant pathogens. PCR-based molecular diagnostics may be able to detect single resistance genes directly from patient specimens more rapidly than any culture-based diagnostics. However, such molecular assays are generally narrow-spectrum assays of single gene targets and also suffer from problems with specificity of resistance genes, resistance that is not genetically-mediated (e.g. upregulation of efflux pumps), and the associated costs^11–13^.

Matrix-Assisted Laser Desorption/Ionization Time-of-Flight (MALDI-TOF) mass spectrometry (MS) has proven to be a rapid technology for microbial species identification. In just a few minutes, MALDI-TOF MS can be used to characterise the protein composition of single bacterial or fungal colonies^14–16^, which are usually available within 24 hours after sample collection. MALDI-TOF MS enables precise and low-cost microbial identification, which has led to the technology becoming the most commonly-used method for microbial identification at species level in clinical microbiology laboratories^7, 17, 18^. MALDI-TOF MS has the potential to move beyond simply identifying an infecting pathogen. Extracting additional information directly from acquired MALDI-TOF mass spectra data may also enable antimicrobial susceptibility testing. Indeed, a recent study used MALDI-TOF mass spectra to detect markers associated with methicillin resistance in clinical samples of *Staphylococcus aureus*^19^. However, the absence of a comprehensive catalogue of marker masses for all potential pathogen and drug combinations has made translating such efforts to clinical practice impossible. In this study, we harnessed the full potential of MALDI-TOF MS to predict antimicrobial resistance through machine learning methods. In this context, previous efforts are rare^20, 21^ and stymied by the lack of large, publicly-available, high-quality benchmark datasets^22, 23^.

To develop clinically-applicable mass spectra-based antimicrobial resistance prediction approaches, we created the **D**atabase of **R**es**I**stance against **A**ntimicrobials with MALDI-TOF **M**ass **S**pectrometry (*DRIAMS)*. DRIAMS is a large-scale, publicly-available, high quality collection of bacterial and fungal MALDI-TOF mass spectra derived from routinely-acquired clinical isolates, coupled with the respective laboratory-confirmed antibiotic resistance profile. We used DRIAMS to undertake the first large-scale study of the utility of such spectra for antimicrobial resistance prediction. We demonstrated the efficacy of this approach for determining the resistance for three priority pathogens, reporting AUROC values of 0.74, 0.74, and 0.80. Furthermore, we validated the ability of DRIAMS to increase resistance profiling performance at other hospitals, where less data is available to build an antimicrobial resistance classifier. We demonstrate the clinical usefulness through a retrospective clinical case study, in which we observe that in 88% of cases, our prediction would have resulted in a beneficial change in the clinical treatment. As mass spectra can be generated rapidly from colonies following overnight culture, our approach can provide guidance for early antimicrobial patient treatment decisions and antibiotic stewardship, substantially sooner than any classical culture-based phenotypic testing.

## Results

### DRIAMS: Clinical routine database combining MALDI-TOF mass spectra and antimicrobial resistance profiles

From 2016 to 2018, we assembled a very large dataset of MALDI-TOF mass spectra from more than 300,000 clinical isolates from four different diagnostic laboratories in Switzerland. The raw dataset comprises a total of 303,195 mass spectra and 768,300 antimicrobial resistance labels and represents 803 different species of bacterial and fungal pathogens. The dataset was processed and organised in four sub-collections (*DRIAMS-A* to *-D*; **Fig. 1**). *DRIAMS-A,* the largest collection with 145,341 mass spectra), was collected at the University Hospital Basel (Switzerland) and is used for the main analysis presented in this study. *DRIAMS-A* contains resistance labels associated with 71 different antimicrobial drugs, the number of spectra and antimicrobial resistance ratios for which can be found in **Suppl. Tab. 1** and **2**. Importantly, the MALDI-TOF mass spectra in DRIAMS-A could be obtained from clinical samples within 24 hours of collection, enabling species identification on a rapid scale as compared to standard phenotypic resistance testing (**Suppl. Fig. 1**).

**Figure 1:**
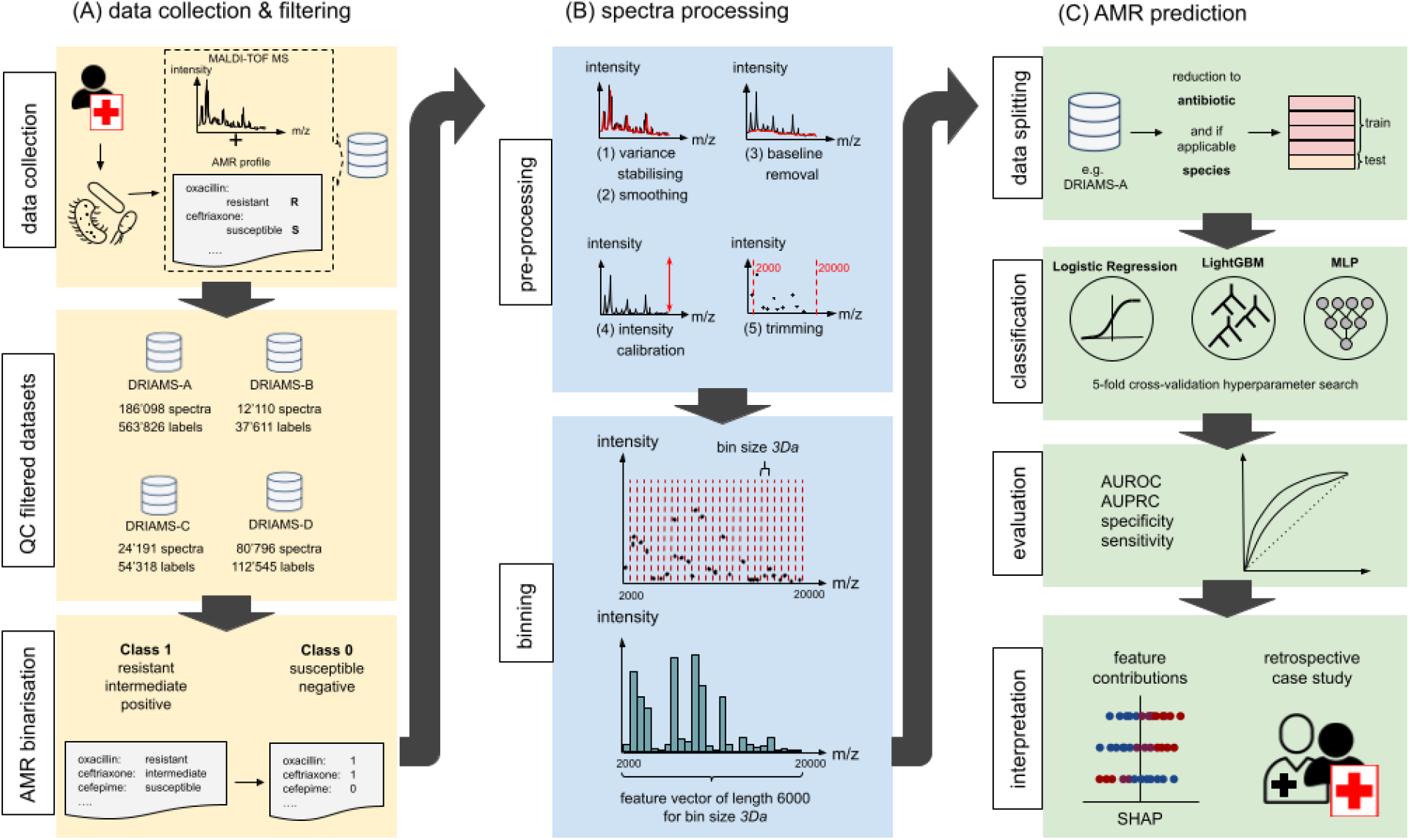
MALDI-TOF MS based AMR prediction workflow. Workflow of MALDI-TOF MS data preprocessing and antibiotic resistance prediction using machine learning. **A.** (i) Data collection: Samples are taken from infected patients, pathogens are cultured, and their mass spectra and resistance profiles are determined. Spectra and resistance are extracted from the MALDI-TOF MS and laboratory information system; corresponding entries are matched and added to a dataset. Samples are filtered according to workstation. (ii) Quality control (QC) filtered datasets: After several quality control steps, the datasets are added to *DRIAMS*. (iii) Antimicrobial resistance (AMR) binarisation: antimicrobial resistance is defined as a binary classification scenario, with the positive class represented by all labels leading to the antimicrobial not being administered, i.e. intermediate or resistant, and positive, while the negative class represents susceptible or negative samples. **B.** (i) Pre-processing: Cleaning of mass spectra. (ii) Binning: Binning spectra into equal-sized feature vectors for machine learning. **C.** (i) Data splitting: For the experiments the samples are subset to only one species. Data is split into 80% training and 20% test, stratified by both antimicrobial class and patient case number. (ii) Classification: Antimicrobial resistance classifiers are trained using a 5-fold cross-validation for hyperparameter search, using the classification algorithms logistic regression, LightGBM, and a deep neural network classifier (MLP). (iii) Evaluation: Predictive performance is measured in metrics commonly used in machine learning (AUROC and AUPRC) and the medical community (specificity and sensitivity). (iv) Interpretation: Interpretation of individual feature contribution to antimicrobial resistance prediction through Shapley values and clinical impact through a retrospective case study on samples from the latest four months of collected data.

### Machine learning for MALDI-TOF MS based antimicrobial susceptibility prediction

To move beyond simple species identification, we preprocessed and binned mass spectra measurement points into fixed bins of 3 Daltons (Da), ranging from 2,000 Da to 20,000 Da, thus obtaining a 6000-dimensional vector representation for each sample. The selected bin size is sufficiently large to adequately represent each spectrum while still remaining computationally tractable (for details see **Methods**). Next, we converted the antimicrobial resistance categories, which are either recorded as *susceptible*, *intermediate*, or *resistant* in the laboratory reports associated with each sample, into a binary label (susceptible vs. intermediate/resistant)(for details see **Methods**). Specifically, we assigned intermediate or resistant samples to the positive class, and susceptible samples to the negative class (in most of the scenarios we consider, the positive class will be the minority class). We then split the samples into training and testing datasets, ensuring that all data associated with a specific case was either part of the train dataset, or the test dataset, but not both, while keeping a similar antimicrobial class ratio in both train and test dataset. We used three machine learning approaches for classification, i.e. logistic regression (LR), gradient-boosted decision trees (LightGBM), and a deep neural network classifier (multi-layer perceptron, MLP), to predict resistance to each individual antimicrobial. The three models were selected because they represent different complexity classes of classifiers (for a more in-depth description of these approaches, please see **Methods**). Subsequently, we report the common machine learning metrics ‘area under the receiver operator characteristic curve’ (AUROC) and ‘area under the precision-recall curve’ (AUPRC) as performance metrics. AUROC can be understood as the probability of correctly classifying a pair of samples, i.e. a resistant/intermediate one and a susceptible one; AUPRC quantifies the ability to correctly detect samples from the smaller of the two classes (resistant/intermediate), while minimising false discoveries. Overall, we observed that LightGBM and MLP were the best-performing classifiers in terms of AUROC. **Fig. 1** depicts the workflow from data collection and filtering, spectra processing, and antimicrobial resistance prediction results.

### Species-specific AMR prediction yields high performance for clinically-relevant pathogens

We first sought to determine whether the use of species-specific mass spectra in *DRIAMS-A* would result in high predictive performance. To this end, we performed a focused analysis for three clinically important pathogens: *Staphylococcus aureus, Escherichia coli*, and *Klebsiella pneumoniae*, all of which are on the World Health Organization ‘priority pathogens’ list^24^. For each of the three species, we selected relevant antibiotics to test based on their clinical usage. We then created a *DRIAMS-A* subset for each antibiotic, which we further divided into stratified training and testing data as described above. For each of the three species we chose one antibiotic resistance as the major scenario of interest; namely oxacillin as a marker for Methicillin-resistant *S. aureus* (MRSA)^25^, and ceftriaxone resistance in *E. coli* and *K. pneumoniae* as marker for extended spectrum or other beta-lactamases (ESBL). We then trained a classifier using each model for each of the three major species-antibiotic pairs (see **Fig. 2A**). We analysed to what extent the respective best model was capable of predicting resistance to other antibiotics (see **Fig. 2B-D**), observing high overall performance in both AUROC and AUPRC values; the classifier is therefore capable of providing precise antimicrobial resistance predictions. For *S. aureus*, the prediction of oxacillin resistance reached a high performance with AUROC of 0.80 and AUPRC of 0.49 at a positive (i.e. resistant/intermediate) class ratio of 10.0%. According to laboratory protocols used in *DRIAMS-A*, for *S. aureus* strains, the reported susceptibility of beta-lactam antibiotics are inferred from the oxacillin susceptibility test results. We also observed high performance for *E. coli* and *K. pneumoniae*, where the prediction of ceftriaxone resistance reached AUROC values of 0.74 in both species, and AUPRC values of 0.30 and 0.33, at a positive class ratio of 10.0% and 8.2%, respectively. We would expect the generation of such resistance information within 24 hours to have a substantial impact in treatment adaptation and infection prevention management. Overall, this experiment demonstrated that a species-specific classifier can achieve clinically useful prediction performance with significantly faster determination of antibiotic resistance compared to the laboratory standard of phenotypic resistance determination (**Suppl. Fig. 1**). We also analysed to what extent the combination of species identity and mass spectrometry information outperforms predictions based on species identity alone. We analysed AUROC predictive performance for the 42 studied antibiotics (see **Suppl. Fig. 2**). For 31 of them, AUROC values above 0.80 were reached, implying highly accurate predictions. Moreover, for 22 antibiotics, we observed statistically significant improvements in prediction performance using the combined mass spectra in *DRIAMS-A* as compared to using only species information for resistance prediction. The results clearly demonstrate the predictive power of mass spectra based antimicrobial resistance prediction.

**Figure 2:**
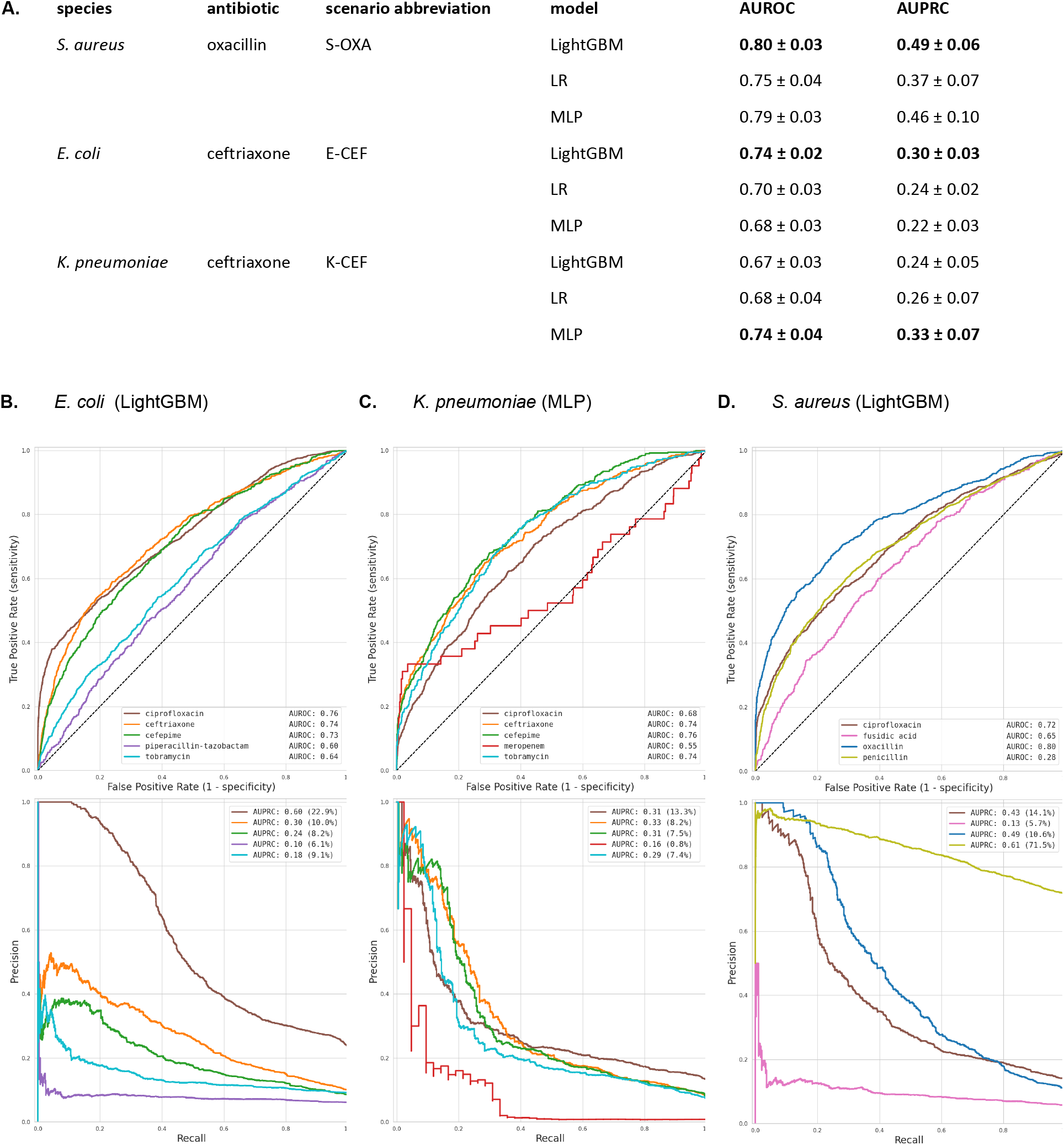
Best performance antimicrobial resistance prediction models on DRIAMS-A. **A. Comparison of performance of three machine learning models.** Metrics are reporting the mean ± standard deviation for 10 different shuffled stratified train–test splits. For *S. aureus* (oxacillin) and *E. coli* (ceftriaxone), the best predictive performance is reached with LightGBM; for *K. pneumoniae* (ceftriaxone) with the multi-layer perceptron (MLP). Additionally, the abbreviations for species-antibiotic scenarios are introduced. **B.-D. ROC and PR curves for different antimicrobials using the best models.** The curves were created by *appending the scores* while the displayed values stem from *reporting the mean* for 10 different shuffled stratified train–test splits, matching values to the tables. **B.** For *E. coli*, the best-performing predictor was that for ciprofloxacin, followed by ceftriaxone, critical antibiotics indicating an extended beta-lactamase (ESBL) if resistant. **C.** For *K. pneumoniae*, cefepime exhibited the highest performance of 0.76 AUROC, also indicating an ESBL if resistant. Compared to the other scenarios, its ROC curve has a larger step size, but with over 500 test samples, the sample size is comparable to the other antibiotics. **D.** Finally, for *S. aureus*, our model performed best for oxacillin, with an AUROC of 0.78. This is particularly relevant, as for *S. aureus*, the resistance to other beta-lactam antibiotics (including amoxicillin/clavulanic acid and ceftriaxone) is directly derived from oxacillin resistance, indicating a methicillin resistant *S. aureus* (MRSA).

### Large external datasets improve local antimicrobial resistance prediction

The use of pre-trained machine learning models could expedite uptake of this approach in clinical laboratories already using MALDI-TOF MS for species identification. As such, we assessed whether predictive performances reached using data from one site (e.g. one specific hospital) are transferable to other sample collection sites. For the datasets *DRIAMS-A* to *-D*, each representing one of our four sites, we divided data associated with each case into train and test datasets as described above, and then trained a predictor before testing on each site. We also compared this site-specific training with predictors trained across all sites. The results indicate that site-specific training reaches better predictive performance compared to across-site validation. Within the site-specific training, the large *DRIAMS-A* dataset is *the* or *among the* best-performing sites (**Fig. 3A**).

**Figure 3:**
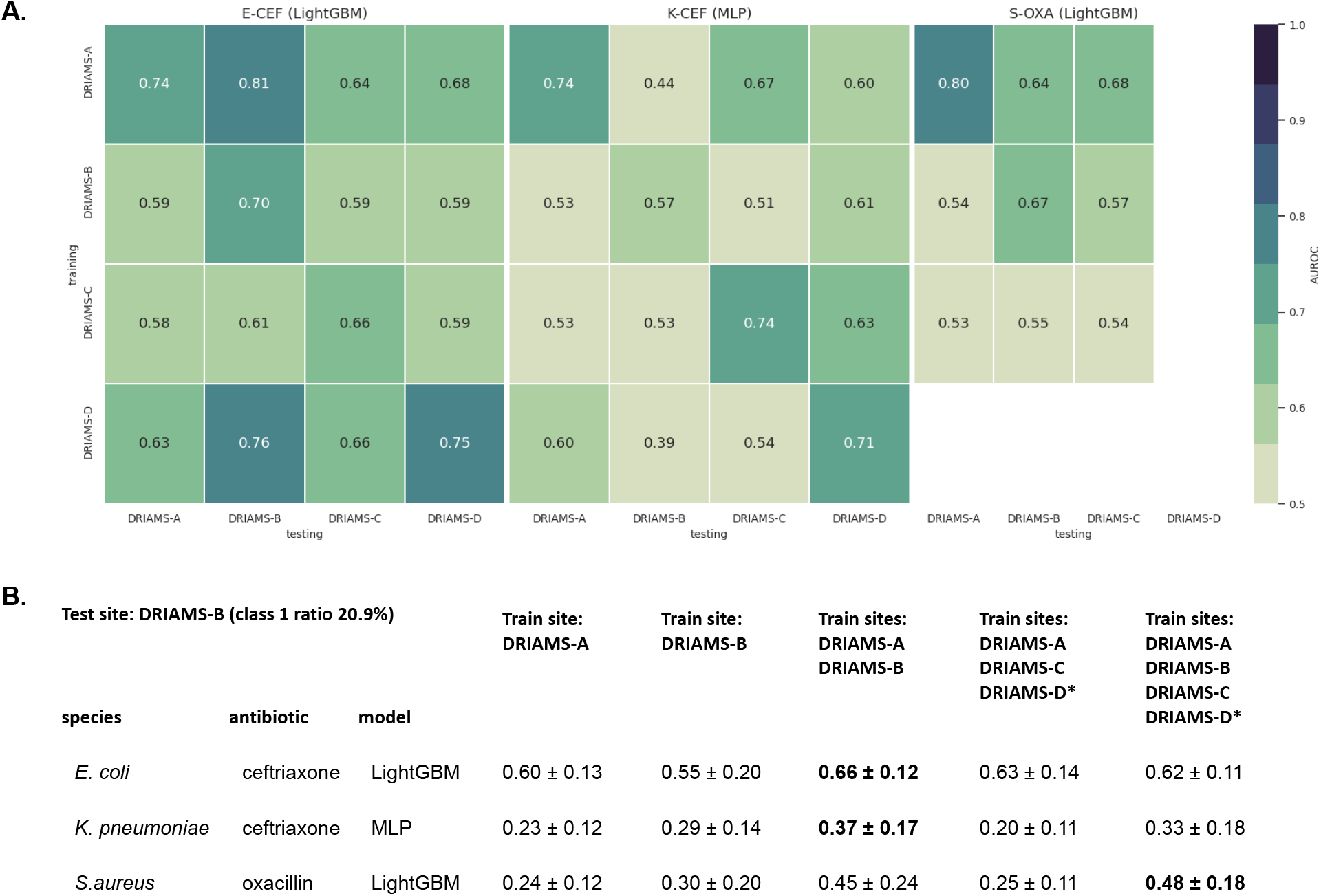
Combining datasets is necessary to reach accurate antimicrobial prediction on external validation sites. **A. Validation predictive performance of each scenario trained and tested on *DRIAMS-A* to *-D* (AUROC).** The results show the mean AUROC performance of 10 random train-test splits. For comparability, the train-test splits are kept the same for each of the respective four *DRIAMS* datasets. The values reported on the top-right (both training and testing *DRIAMS-A*) correspond to the values reported in **Fig. 2A**. With the exception of *DRIAMS-B E.coli* (ceftriaxone), the highest performance is reached when training is performed on the same site as testing. *DRIAMS-A* and *DRIAMS-D* exhibit the highest transferability with respect to predictive performance, and overall, transferability seems higher in *E. coli* as compared to *K. pneumoniae* and *S. aureus*. Due to the different class ratios between test datasets on different sites, AUROC was chosen to permit comparability. The scenario abbreviations follow **Fig. 2A**. For S-OXA no *DRIAMS-D* data is available. **B. Performance trained on union datasets of multiple sites and tested on *DRIAMS-B* (AUPRC).** For each site the dataset was split into training and test; only the training sets are combined in the union training sets. Neither the large external dataset DRIAMS-A nor the internal datasets at the target site alone reach the best performance on the target site test data. A union of the target training data and large external datasets is able to reach significant improvements over the target site performance. (*) For S-OXA no *DRIAMS-D* data is available. The results for test sites *DRIAMS-C* and *DRIAMS-D* are listed in **Suppl. Tab. 3**.

We further investigated whether we could improve prediction for sites where a large dataset is unavailable by leveraging existing large external datasets such as *DRIAMS-A*. We therefore trained on combinations of training datasets from different sites, including different combinations of the four sites *DRIAMS-A* to *-D*, and tested on a single external site *DRIAMS-B* to *-D*. While the transferability of predictive performance from one site to another is an active area of research in the machine learning field of domain adaptation, a recent study^26^ has shown that using empirical risk minimization by learning a single model on pooled data across all training environments often outperforms more complex domain adaptation approaches. The results indicated that the addition of training datasets from other sites to the external site train data was beneficial for validation sites *DRIAMS-B* and *-C* (**Fig. 3B** and **Suppl. Tab. 3**). For external validation site *DRIAMS-D*, the best predictive performance was still reached when training exclusively on the site-specific training data. For external validation sites *DRIAMS*-B, the addition of the large *DRIAMS-A* dataset proved most beneficial for scenarios *E. coli* (ceftriaxone) and *K. pneumoniae* (ceftriaxone), while adding more data from *DRIAMS-C* to the training data was beneficial for *S. aureus* (oxacillin).

### Learning on a single species yields superior predictions compared to learning on multiple species

Next, we analysed whether classifiers can improve the predictive performance by training on a large number of samples from multiple species (as opposed to training on samples from a single species). It is known that different species of bacteria can be resistant to a specific antimicrobial through different mechanisms. For example, resistance against beta-lactam antibiotics in Gram-negative bacteria, such as *E. coli*, may be caused by the production of beta-lactamases such as CTX-M^27^, TEM, and SHV^28, 29^ or carbapenemases e.g. OXA-48^30^. Resistance against beta-lactam antibiotics in Gram-positive bacteria, such as *S. aureus*, can be caused by a penicillinase (blaZ), resulting in a resistance only against penicillin^31^, or by an alteration within the penicillin-binding protein (PBP2a), resulting e.g. in the MRSA phenotype with resistance against multiple beta-lactam antibiotics^32^. Hence, pooling spectra across species and predicting antimicrobial resistance using the same model regardless of the species poses a more complex learning task than predicting antimicrobial resistance within one specific species. However, stratification by species reduces the number of samples available for training and might therefore lower predictive performance. We assessed the trade-off between the number of available samples and predictive performance by comparing the performance of (i) a model trained to predict antimicrobial resistance using samples from across all species (*ensemble*) with (ii) a collection of models trained separately for *single* bacterial species (**Fig. 4A**). Each point of the depicted curves corresponds to one classifier, trained with the number of samples specified on the x-axis. The last, i.e. rightmost, point of each curve hence corresponds to the scenario in which all available samples are being used. We observed that training a model for individual species *separately* led to improved performance for all species, despite the reduction in sample size. Notably, all training samples used to reach the last single-species classification results were also included in the training samples for the last ensemble classifier. The last ensemble classifier therefore had access to at least the same amount of information about the respective species as the last single-species classifier. Nevertheless, it never outperformed the single-species classifiers except for oxacillin resistance in *S. aureus*. Furthermore, a few curves reached a plateau, with the single-species classifier increasing more sharply with the last addition of more training samples. This demonstrates the higher complexity of the ensemble prediction task and the benefit of a larger training dataset, which are critical for capturing different resistance mechanisms.

**Figure 4:**
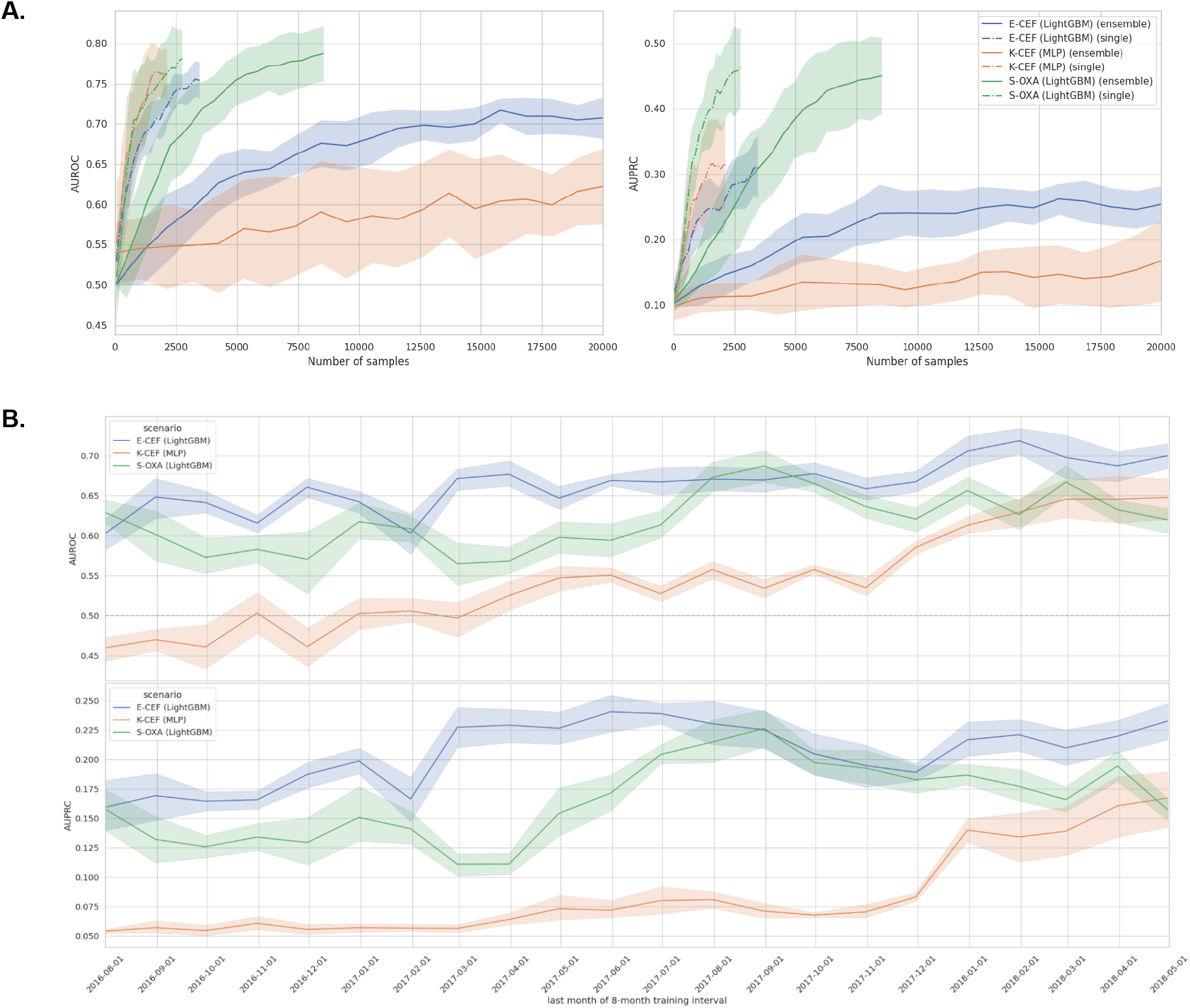
Stability of results with different dataset perturbations. The scenario abbreviations follow Fig. 2A. **A. Predictive performance with increasing sample size** AUROC and AUPRC as a function of sample size for complete and species-stratified *DRIAMS-A* datasets. Experiments were repeated for ten different shuffled train–test splits. The solid curve represents the mean of these repetitions, with the envelope depicting the standard deviation of repetitions. Results are shown for the three major scenarios of interest. With equal sample size, training only on samples from a single species is outperforming training in all scenarios. Even for the datasets containing all available samples from the target species (the rightmost points of each curve), the single-species scenario outperforms the ensemble in both *E. coli* and *K. pneumoniae* (ceftriaxone), while the curves reach a similar predictive performance for *S. aureus* (oxacillin). **B. Temporal validation in *DRIAMS-A* reporting AUROC (upper) and AUPRC (lower) of sliding 8-month training window on fixed test set.** The test dataset is the data collected in the last four months May to the end of August 2018. For *E. coli* (ceftriaxone) (E-CEF) and *S. aureus* (oxacillin) (S-OXA) the predictive performance decreases with increasing temporal distance to the test set, but the fluctuations in the curve are of the same size as the drop over time. The predictive performance for ceftriaxone in *K. pneumoniae* (K-CEF) decreases more continuously and drastically with increasing temporal distance to the test set.

### Current samples are advantageous for accurate antimicrobial resistance prediction

Mass spectra profiles are subject to variations and differences over time, caused by biological differences through the ongoing evolution of the local microbial populations (with new strains being introduced by e.g. travelling), or by technical differences, such as changes after MALDI-TOF MS machine maintenance (e.g. laser replacement and adjustment of internal spectra processing parameters through machine calibration). To guide and encourage further method development, we wanted to illustrate challenges and limits of mass spectra based antimicrobial resistance prediction. Hence, we studied whether recent samples are necessary, and whether adding more samples collected at older timepoints would increase predictive performance. We fixed the latest four months of data from *DRIAMS-A* as a test dataset, and trained classifiers on data collected within 8-month training windows with increasing temporal distance to the test collection window, simulating the availability of older samples. The training data within the training window was oversampled to match the class ratio in the test data; however, sample sizes could still vary between training windows. We observed a slight *decrease* in performance with increasing temporal distance between training and testing data (**Fig. 4B**) for *E. coli* and *S. aureus;* with a larger decrease for *K. pneumoniae*. We explain this drop by the aforementioned differences that accumulate over time, highlighting the positive effect of having access to *recent* training samples.

### Retrospective clinical case study

In order to evaluate the clinical benefit of our classifier, we evaluated the antibiotic therapy of patients represented in *DRIAMS-A,* with invasive serious bacterial infections treated between April and August 2018. We reviewed 416 clinical cases that included positive cultures with *E. coli*, *K. pneumoniae,* or *S. aureus* from either blood culture or deep tissue samples. For 63 of these cases, an infectious diseases specialist (hereafter referred to as *clinician*) was consulted regarding the antibiotic treatment. Consultation occurred between the species being identified and before the phenotypic antibiotic resistance testing was available (**Suppl. Fig. 3**). For each case, we retrospectively reviewed the recommendations and assessed whether an alternative antibiotic therapy would have been suggested if our classifier had been employed at the time at which the MALDI-TOF mass spectrum was acquired.

In 54 clinical cases, the employment of the algorithm would not have changed the suggested antibiotic treatment: in 22 cases the clinician suggested de-escalation of the antibiotic regimen to a more-narrow spectrum antibiotic, in 25 cases to continue the current antibiotic regimen, and in seven cases to escalate the antibiotic treatment to a broader spectrum antibiotic. The classifier reported an accurate prediction of the antibiotic resistance in 51 of these 54 cases, but as the decision on antibiotic treatment can be influenced by multiple factors other than the antibiotic resistance of one bacterial species against one antibiotic agent, such as allergy, these did not change the suggested therapy (**Fig. 5**). In three cases, our algorithm predicted susceptibility, where phenotypic testing revealed resistance to antibiotics. In none of these three cases however, would this incorrect prediction have led to a less effective treatment than suggested without the algorithm: In two of these cases a known MRSA colonization of the patient would have been considered by the clinician, regardless of the prediction of the algorithm. In the third case, *K. pneumoniae* and *E. coli* were both identified in blood culture samples. Here, the clinician would have suggested to keep the current antibiotic treatment against *E. coli with or without the use of the algorithm,* and escalation to a broader spectrum antibiotic was only implemented after phenotypic testing.

**Figure 5:**
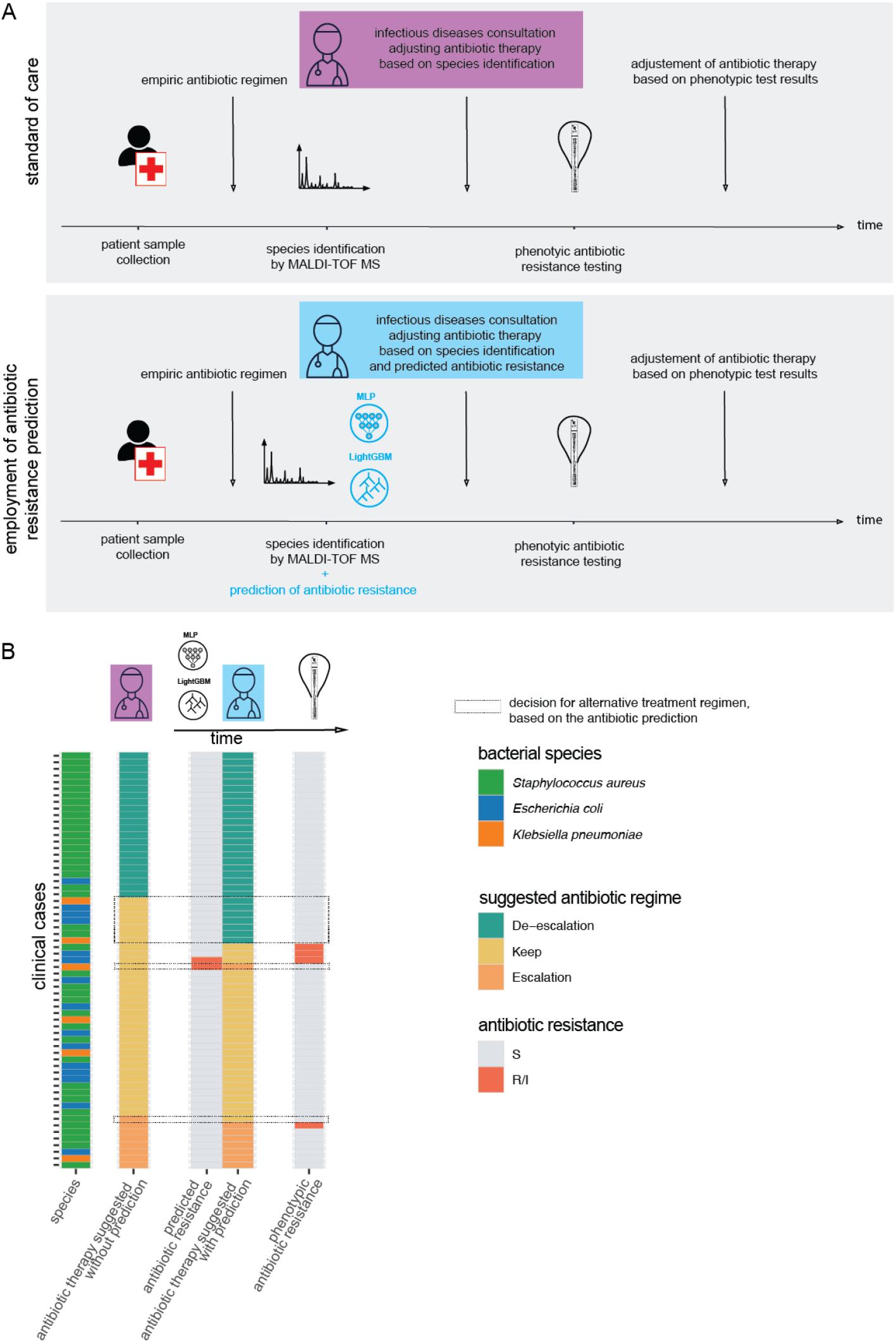
Retrospective clinical case study including 63 cases with invasive bacterial infection. (A) schematic representation of the current standard of care (top row) and the possible employment of our classifiers in the clinical workflow (bottom row). (B) We evaluated the antibiotic regime suggested by a clinician without the employment of the classifier (column 2), which antibiotic resistance the classifier predicted (column 3), the antibiotic treatment suggested considering the predicted antibiotic resistance (column 4) and the phenotypically tested antibiotic resistance. The dashed boxes highlight where the employment of the classifiers would have led to an alternative antibiotic treatment suggestion. ‘de-escalation’: change antibiotic regimen to a more narrow spectrum antibiotic agent, ‘keep’: continue the current antibiotic regimen, ‘escalate’: change antibiotic regime to a broader antibiotic regimen.

For nine cases an alternative antibiotic therapy would have been suggested by the clinician with the employment of the classifier at the time of species identification: in seven cases, the classifier would have led to a de-escalation of the antibiotic therapy, while in one case, the employment of the algorithm would have led to an escalation of the antibiotic therapy, and finally, in one case the employment of the algorithm would have changed the suggested treatment to continue the current antibiotic therapy, where the clinician suggested to escalate to an antibiotic agent with a broader spectrum (**Fig. 5**). In summary, for eight out of these nine cases (88%) where the employment of the algorithm would have changed the empiric antibiotic regimen, this change would have been beneficial and would have promoted antibiotic stewardship.

### Analysis of feature contributions through Shapley values

Only very few studies have considered full mass spectrum information instead of single peaks for antimicrobial phenotype prediction^20^. We therefore wanted to assess whether predictive performance is primarily driven by only a subset of the peaks, or whether the full spectrum is employed. While this question is partially addressed by the use of feature importance values, their use without additional information can be misleading as their interpretation is highly contingent on the classifier that was employed. Hence, for further analysis, we also calculate the Shapley values, a concept originating from coalitional game theory, which enable the interpretation of model output contributions on both the dataset and per-sample level for each feature^33^. **Fig. 6** visualizes the average and per-datapoint Shapley values for the 30 features with the highest average contribution. It is evident that three to ten mass-to-charge bins contribute more than the remaining features. As the tails of the distribution plots for each feature are coloured with either the highest or lowest feature value, we see that the predictor is using either the presence of a high intensity value (red) or the absence of any measured intensity (blue) for a positive (resistant/intermediate) class prediction. In case of *S. aureus* (oxacillin) for the top four mass-to-charge bins the presence of a peak indicates the positive (resistant/intermediate) class, while for *E. coli* (ceftriaxone) also the absence of a peak can strongly contribute to a positive class prediction. We further observe that most of the top 30 contributing features lie in the lower mass-to-charge ratio regime (lower than 10,000 Da), where more mass particles are measured in MALDI-TOF MS. The feature importance distributions over all 6,000 features (**Suppl. Fig. 4**) stemming directly from the classification models indicate that the classifiers utilize the entire range of features.

**Figure 6:**
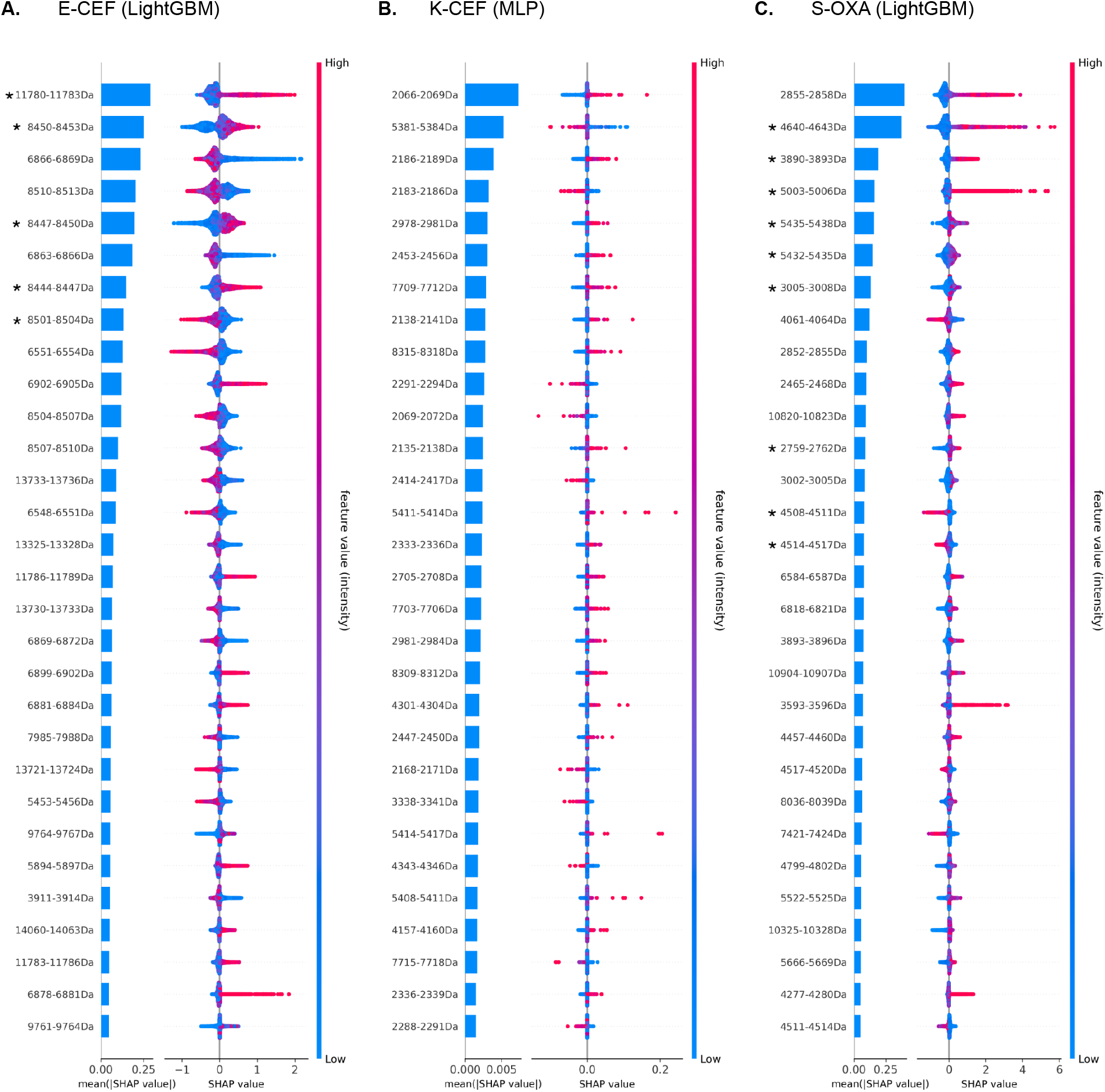
Quantification of feature impact on prediction through analysis of SHapley Additive exPlanations (SHAP) values of the 30 most impactful features. For each scenario, a barplot on the left indicates the mean Shapley value, i.e. the average impact of each feature on the model output magnitude. The scatterplot on the right indicates the distribution of Shapley values, and their impact on the model output, over all test samples. The colours of each test spectrum (according to the colorbar: blue for low feature values and red for high feature values) indicates the feature value, i.e. the intensity value of the respective feature in the spectrum. The scenario abbreviations follow Fig. 2A. The asterisks marked feature bins containing a previously identified protein peak listed in **Suppl. Tab. 4**.

## Discussion

We have demonstrated that MALDI-TOF mass spectra based antimicrobial resistance prediction from routine diagnostic clinical samples is capable of providing accurate predictions with unmatched speed within 24 hours of sample collection. This analysis was made possible by collecting the largest real-world clinical dataset containing 303,195 MALDI-TOF mass spectra and 768,300 corresponding antimicrobial resistance phenotypes. Overall, we observed high predictive performance using calibrated LightGBM and MLP classifiers for multiple pathogen species with classifiers trained on individual species–antibiotic combinations, such as ceftriaxone resistance in *E. coli* and *K. pneumoniae* and oxacillin resistance in *S. aureus*, obtaining AUROC values larger than 0.70 for several antibiotic–species scenarios (**Fig.2**).

We found that antimicrobial resistance classifiers trained on mass spectra of species from one site are not directly applicable to mass spectra measured at other sites (see **Fig. 3A**). This is influenced by many sources, including (i) different phylogenetic strains, (ii) different prevalence of resistance (i.e. different class ratios), which can impact predictive performance, or (iii) technical variability^34^, owing to different machine-specific parameters and settings (i.e. ‘batch effects’). In a similar vein, the closer the time of collecting the training samples is to the time of prediction, the better the predictive power of the trained classifier (see **Fig. 4B**), likely owing to the same aforementioned reasons. Hence, we would recommend that a clinically-applied classifier should be updated and retrained with the most recent data, originating from its deployment site, at regular time intervals. In clinical practice such an algorithm may require regular re-certification. At the same time, for individual specific species–antibiotic scenarios, even small sample sizes can lead to high predictive performance (see **Fig. 4A** and the subsequent discussion), making it possible to train hybrid models based on e.g. *DRIAMS-A* with few site-specific examples.

We demonstrate that in order to obtain a classifier at a site with a smaller training dataset, combining the available data with an external dataset, such as *DRIAMS-A,* can increase the training performance (**Fig. 3B** and **Suppl. Tab. 3A**). Combining training datasets from different sites increases the sample size, and potentially the coverage of rarer bacterial strains, which improves the predictive performance. However, combining training data originating from different sites also increases the variance in the data, which has the potential to decrease predictive performance. Merging training datasets did not lead to an increased performance on the *DRIAMS-D* test data (**Suppl. Tab. 3B**). This could indicate that *DRIAMS-D*’s outpatient sample pool creates a dataset dissimilar to the dataset collected at hospitals (*DRIAMS-A* to *-C*). These results motivate the potential of large-scale MALDI-TOF MS clinical routine dataset acquisition for antimicrobial resistance prediction—combining large datasets could increase the predictive performance on either prediction site. Furthermore, it is worth noticing that all collection sites contributing data to this study are located in relatively close geographic proximity in a low endemic area for ESBL producing bacteria. Future analysis should assess how data from healthcare centers with a higher burden of antibiotic resistant bacteria influence the performance of our classifiers.

Surprisingly, we found the predictive performance of classifiers trained on a single species to be higher than that of classifiers trained on multiple species, indicating the higher complexity of predicting multiple resistance mechanisms. However, we also observed the general trend of improved performance if more samples are available. This indicates the potential benefits of having access to a large database of MALDI-TOF mass spectra (**Fig.4**). **Suppl. Fig. 5** also indicates that the reduction in performance training on older datasets could in fact stem from a lower sample size, as the MALDI-TOF MS technology usage at the *DRIAMS-A* collection site increased over time. **Fig. 4A** demonstrates that classifiers perform best for a specific species–antibiotic scenario if the training dataset includes only samples from that scenario, and that adding samples from other species lowers the predictive performance. We distinguish two potential situations: (i) the mechanism causing antimicrobial resistance varies between species or (ii) the resistance mechanism is the same in several species. If situation (i) were true, the results in **Fig. 4A** can be expected, as the classifier has to predict several objectives at once (predicting resistance caused by mechanism A in species A, while also being able to predict resistance caused by mechanism B in species B, etc.). In situation (ii), one could expect that combining samples of several species to predict resistance caused by the same mechanism increases performance. For most scenarios, we are confronted with situation (ii), such as beta-lactam resistance in *E. coli* in **Fig. 4A**. However, many proteins causing resistance are beyond the effective mass range of MALDI-TOF mass spectra. For example, the penicillin-binding protein in *S. aureus* has a mass of approximately 76,400 Da^35^, beta-lactamases in *E. coli* and *K. pneumoniae* weigh approximately 30,000 Da^36–39^, and the *E. coli* outer membrane porin OmpC weighs approx. 40,300 Da^40^. Therefore, we hypothesise that our predictor cannot detect the resistance mechanism (due to physical limitations), but rather species-specific and resistance-associated changes in the proteome as well as phylogenetic similarity between resistant vs. susceptible samples. The number of samples depicted in **Fig. 4A** also gives an indication at what sample size most information/variance of samples originating from the *DRIAMS-A* collection site are covered by the training data. The sample sizes required to reach the ‘plateau’ visible for single-species predictive performance range from 2,500 to 5,000. We therefore suggest collecting a dataset of at least 2,500 samples when working on MALDI-TOF MS based antimicrobial resistance prediction.

While the antimicrobial resistance classifiers are trained and predict resistance labels as a black-box system, analysing the contribution of each feature bin to the predictive outcome is of utmost importance to explain the antimicrobial resistance predictor decision-making process in a manner that can be interpreted by the user. We therefore determined the feature importance (given by the respective algorithms LightGBM and the MLP) and the Shapley values of each feature bin and compared the results of the highest-weighted bins to known resistance associated peaks from the literature. We first note that most of the feature bins with the highest average impact are feature bins with a mass-to-charge of less than 10,000 Da (79 out of 90 features bins in **Fig. 6**) and the Shapley values indicate that very high or very low feature bin values (corresponding to the presence or absence of a MALDI-TOF mass peak within the feature bin range) contribute to the prediction outcome, rather than variations in the feature bin magnitude. This is in line with prior knowledge on MALDI-TOF MS; most proteins that are reproducibly detected in MALDI-TOF MS have a weight less than 10,000 Da^41^ and the signal indicates their presence or absence. This confirms that the detection of proteins is responsible for the predictive power, rather than confounding signals or noise.

Multiple feature bins that contributed substantially to our classifiers can be annotated with resistance-associated proteins identified in previous studies (**Suppl. Tab. 4**). Most studies aiming to identify resistant bacterial strains from routinely acquired MALDI-TOF mass spectra have focused on oxacillin resistance in *S. aureus* and have identified peaks that were either used to distinguish between methicillin susceptible *S. aureus* (MSSA) and MRSA or to distinguish between MRSA sub-lineages^42–50^. A subset of these discriminatory peaks were identified to correspond to either (i) constitutively conserved housekeeping genes or (ii) other peptides such as stress proteins or low molecular weight toxins^44^. The mass of three of the identified proteins, 3007 Da (Delta-toxin), 3891 Da (uncharacterised protein SA2420.1), and 4511 Da (uncharacterised protein SAR1012) can be attributed to highly contributing feature bins; specifically 3005-3008 m/z, 3890-3893 m/z, and 4508-4511 m/z receiving the 7th, 3rd, and 14th Shapley value (**Fig. 6, Suppl. Tab. 4**) respectively. A peak at 2415 m/z has previously been identified as MRSA specific^43^. This peak corresponds to the peptide PSM-mec^46^, which is encoded on a subset of *SCCmec* cassettes (types II, III and VIII) in close proximity to *mecA^51^*^52^, which encodes resistance to oxacillin. Among the feature importances within the oxacillin resistance predictor in *S. aureus*, this peak corresponds to the 83rd highest-ranked feature bin of 2414-2417 m/z (out of 6000 feature bins overall).

The increased occurrence of multidrug resistant *E. coli* has been attributed to the spread of a few clonal lineages, in particular to sequence type (ST) 131^53^. Previous studies^54^ have identified ST131 characteristic peaks, which can be attributed to feature bins receiving high feature importances and Shapley values by the ceftriaxone of *E. coli* (8447-8450m/z, 8498-8501m/z and 11780-11783m/z; receiving the 5th, 38th, 1st highest feature importance respectively). These references confirm the discriminatory potential of single feature bins, contributing substantially to our classifiers and also highlight their generalisability, as the spectra for these studies were acquired from independent strain collections and on different MALDI-TOF MS devices. Moreover, our classifiers use many more feature bins, for which the discriminatory potential has not previously been identified. An investigation of the protein identity of these yet unknown discriminatory feature bins and their occurrence throughout the respective species would be desirable in the future.

Our retrospective clinical case study shows that our classifier can have a beneficial impact on patient treatment and promote antibiotic stewardship. In 51 out of 63 cases, employing the algorithm at the time of the species identification would have supported the treatment regimen suggested by the clinician (**Fig. 5**). In three cases, the inaccurate prediction by our classifier would not have changed the suggested antibiotic regime, since the decision is influenced by multiple other factors in addition to the resistance profile towards one antibiotic such as (i) allergies of the patient, (ii) other bacterial species involved in the infection, (iii) patient history including the antibiotic profile of previous isolates, (iv) type of administration of the antibiotic agent. In eight out of 63 cases the accurate prediction by our algorithm would have led to an earlier streamlining of the antibiotic regimen to a more narrow spectrum antibiotic agent, which would have been an important improvement considering the urgent threat to public health posed by the spread of antibiotic resistant bacteria. Similar trends in antibiotic stewardship have been observed when using genotypic assays such as rapid PCR assays^55^. These findings exemplify the potential of classifiers to optimize antibiotic treatment and assist antibiotic stewardship efforts using real clinical cases. The evaluation of our classifier in prospective clinical studies, on multiple sites with different prevalence of antimicrobial resistant bacteria, will be necessary to fully evaluate its clinical impact. Clearly, the prediction of resistance alone would not be used, but the prediction may support clinical decision-making that also considers additional patient-related factors.

In summary, our work demonstrates that MALDI-TOF MS based machine learning can provide novel ways to predict antimicrobial resistance in clinically highly-relevant scenarios. The results demonstrate the benefit of large sample sizes on predictive performance. Further work could build upon these findings and leverage unlabeled (no antimicrobial resistance profile available) MALDI-TOF mass spectra in DRIAMS for pre-training a classifier in a semi-supervised fashion before fine-tuning the model on the labeled dataset. In addition to potentially improving the prediction performance, such a training setup could result in a transfer learning scenario to mitigate batch-effects between different collection sites. While these idiosyncratic challenges need to be overcome, there is also a large potential to improve patient treatment.

## Online Methods

### Reproducibility of Results and Data availability

All R and Python scripts can be found in https://github.com/BorgwardtLab/maldi_amr.

### MALDI-TOF mass spectra acquisition and antimicrobial resistance testing

We collected data from daily clinical routine at ISO/IEC 17025 accredited diagnostic routine laboratories. The study was evaluated by the local ethical committee (IEC 2019-00729). Specifically, all MALDI-TOF mass spectra contained in *DRIAMS-A* to *-F* were acquired at four microbiological laboratories in Switzerland providing routine diagnostic services for hospitals and private practices. All laboratories use the Microflex Biotyper System by Bruker Daltonics (Bremen, Germany), which is a widely-employed MALDI-TOF MS system in microbiological routine diagnostics both in North America^16^ and in Europe^17, 18^. The four diagnostic laboratories included in this study are (1) University Hospital Basel-Stadt (providing *DRIAMS-A*), (2) Canton Hospital Basel-Land (providing *DRIAMS-B*), (3) Canton Hospital Aarau (providing *DRIAMS-C*), and (4) laboratory service provider Viollier (providing *DRIAMS-D*). While Canton Hospitals Basel-Land and Aarau employ the Microflex Biotyper LT/SH System, Viollier uses the Microflex smart LS System. Although these two systems differ in their respective laser gas, they use the same reference spectra database, so we included spectra of both Microflex Biotyper systems. University Hospital Basel-Stadt uses the two Microflex Biotyper systems in parallel. The species of each mass spectrum was identified using the Microflex Biotyper Database (MBT 7854 MSP Library, BDAL V8.0.0.0_7311-7854 (RUO)) included in the flexControl Software (Bruker Daltonics flexControl v.3.4). Similar to the mass spectra, antimicrobial resistance profiles were routinely acquired in the same four microbiological laboratories within the same time frames of the dataset. Resistance categories for bacteria values were determined either using microdilution assays (VITEK® 2, BioMérieux, Marcy-l’Étoile, France), or by minimal inhibitory concentration (MIC) stripe tests (Liofilchem, Roseto degli Abruzzi, Italy), or disc diffusion tests (ThermoFisher Scientific, Waltham, USA). Resistance categories for yeast were determined by using Sensititre Yeast One (Thermofisher). All breakpoint measurements were interpreted to be either susceptible, intermediate, or resistant according to EUCAST ^56^ and CLSI (2015 M45; 2017 M60) recommendations. The EUCAST versions used were updated with every EUCAST Breakpoints table update and include v6-v8.

### Quality control

Empty spectra and calibration spectra were excluded from further analysis. This serves to ensure a similar level of data quality for the different sites.

### Matching of MALDI-TOF mass spectra and antimicrobial resistance profiles

MALDI-TOF MS based antimicrobial resistance prediction requires a dataset containing mass spectra and their corresponding resistance labels, in the form of antimicrobial resistance profiles. In order to construct such a dataset, MALDI-TOF MS and resistance profile measurements belonging to the same microbial isolate have to be matched. Since each site in *DRIAMS* stores the mass spectra and their corresponding antimicrobial resistance profiles in separate databases, a matching procedure has to be developed for each site.

We use the term ‘laboratory report’ for the document used to report laboratory measurement results, including antimicrobial resistance profiles, for each patient within the clinical care. The species of the specimen is obtained through Bruker Microflex MALDI-TOF MS and added to the laboratory report. This decouples laboratory report entry and the mass spectrum; there is no link required between the spectrum file and the laboratory entry after the species is entered. The antimicrobial resistance profiles obtained in their individual experiments are also added to the laboratory report. The laboratory report entries are commonly identified by codes linking them to a patient, or a unique sample taken from a patient, to which we refer as ‘sample ID’. Multiple entries with the same sample ID can exist if several probes were taken from the same patient or several colonies tested from the same probe.

In general, the spectra recorded by the Bruker Microflex systems were labelled with an ambiguous, i.e. non-unique, code corresponding to the non-unique sample ID in the laboratory report. MALDI-TOF mass spectra and their corresponding antimicrobial resistance profiles were stored in separate files. In the clinic, MALDI-TOF MS spectra are never intended to be matched up with the laboratory report entries; therefore no proper protocols for matching exist. Matching protocols had to be developed uniquely and in an ad-hoc fashion for each labelling system at each institution.

In order to link mass spectra to their antimicrobial resistance profiles, we constructed a *unique* identifier, using the sample ID and the determined genus of a sample. The rationale behind this strategy is that if multiple sample ID entries exist, this is most likely due to multiple genera being present in the patient samples, leading to several measurements. We omitted samples for which we were unable to construct a unique sample ID–genus pair. Mass spectra were stored without information on the determined species. Hence, for each spectrum, the species and genus label is determined by re-analysing the spectra with the University Hospital Basel-Stadt Bruker library and then matching the spectrum to its corresponding antimicrobial resistance profile using the assigned sample ID and the determined genus. All MALDI-TOF MS systems used in this study were maintained according to the manufacturer’s standard and spectra were routinely acquired using the ‘AutoXecute’ acquisition mode. The genus is used (instead of species), as it allows for some flexibility between the species assigned to a sample in the laboratory report and the Microflex Biotyper. The species label given in the laboratory report can differ from the species assigned to the corresponding MALDI-TOF mass spectrum by the Microflex Biotyper System as additional microbiological tests can give a more accurate label. In what follows, we provide additional details regarding the matching procedure which are specific to each site.

#### University Hospital Basel-Stadt

Starting in 2015, the spectra were labelled with a 36-position code by the Bruker machine (e.g. 022b130c-6c8c-49b5-814d-c1ea8b2e7f93), which we term ‘Bruker ID’. This code is guaranteed to be unique for all spectra labelled from one machine. Each AMR profile is labelled with a 6-digit sample ID, which is unique for samples in one year. Antimicrobial resistance profiles were collected using the laboratory information system. The laboratory information system includes all entries made for a sample, also entries which have later been corrected and have not been reported nor considered for patient treatment. As such manual corrections are very rare, the uncertainty in antimicrobial resistance labels is limited. For each year (2015, 2016, 2017, and 2018) there are separate antimicrobial resistance profile tables and folders containing all spectrum samples collected during the corresponding year. We lost 40,569 spectra out of 186,098 by following the aforementioned pre-processing routines (*DRIAMS-A*).

#### Canton Hospital Basel-Land

The antimicrobial resistance profiles and mass spectra are each labelled with a 6-digit sample ID. The genus depicted in each mass spectrum was determined through comparison to the Microflex Biotyper Database (Bruker Daltonics flexControl v.3.4); the genus of each antimicrobial resistance profile was stated in the laboratory report. Mass spectra and antimicrobial resistance profiles were merged using the 6-digit sample ID and the genus information.

#### Canton Hospital Aarau

Here, the laboratory report contains the 10-digit sample ID, species label, and antimicrobial resistance profiles of measured samples. This software version did not provide a unique 36-character code for each spectrum, but only a 10-digit sample ID that had to be used to match spectra to the antimicrobial resistance profiles from the laboratory. Since the sample ID can be shared by different spectra, it cannot be used to uniquely match a species label to an input spectrum. To circumvent this problem, we divided the spectra in 15 batches, each one only containing unique 10-digit sample IDs. Repeated sample IDs were distributed over the batches. These 15 batches were re-analysed and labelled by the Bruker software, and 15 output files with the given species labelled were created. Through the separation in batches, the certain species label was determined for each spectrum. The label for each spectrum in the batches can be determined, as we only included spectra that already had a label in the lab file. Now, each spectrum file has a combined label made up of its 10-digit sample ID and its species label. If this combined label was found to have a unique match within the lab results file, the AMR profile was assigned to the spectrum, otherwise its antimicrobial resistance profile position remained empty and only the spectrum with its species label was added to the dataset. We ignore all spectra that could not be matched to an entry in the lab results file (such spectra arise from measurements that do not provide AMR information).

#### Viollier

While all other sites reported AMR labels with either ‘R’, ‘S’, ‘I’, ‘positive’ or ‘negative’ values, samples provided by Viollier are labelled with precursory measurements, namely the MIC of each antibiotic. We therefore use the breakpoints given the up-to-date EUCAST guidelines (v.9) to convert the MIC values to ‘RSI’ values. 80,796 spectra in the `fid` file format are present, identified again through a unique 36-character ‘Bruker ID’. The antimicrobial resistance results are identified by a 10-digit sample ID, which are linked to the Bruker IDs in an additional file, the ‘linking file’. The main reasons for loss of data in preprocessing are (1) the antimicrobial resistance results and ID ‘linking files’ contained significantly fewer entries than `fid` files present (40,571 and 51,177 respectively) and (2) following advice by the lab personnel, only the 10-digit sample ID could be used for matching to the BrukerID (which contained a longer version of the LabID). Through exclusion of all entries without a unique 10-digit sample ID in both the antimicrobial resistance results and Iinking files, another significant portion of data was lost. Specifically, there is an overlap of 10,852 filtered entries from the laboratory report file and the linking file. After matching these entries with spectra, 7,771 spectra with 7,720 antimicrobial resistance profiles remained. Spectra without an antimicrobial resistance profile are not used for any supervised learning tasks (such as prediction).

### Conversion to *DRIAMS*

We require a DRIAMS dataset entry to contain (1) a spectra code linking unambiguously to a spectra file, (2) a readable mass spectra `fid` file (as all of our mass spectra are measured on Bruker Microflex machines), (3) the corresponding species (species reported as ‘Organism best match’ by the flexControl Software was assigned), and, if available, (4) AMR profiles stating antimicrobial susceptibility results.

### Workstations

Patient samples processed at the USB microbiological diagnostic laboratory samples are analysed at nine different workstations, split by patient isolation material or procedure. The workstations comprise (i) urine samples, (ii) blood culture samples, (iii) stool samples, (iv) genital tract samples, (v) samples for which a PCR-based test were ordered, (vi) respiratory tract samples, (vii) samples from deep, usually sterile material (viii) samples that were collected for the hospital hygiene department, and (ix) samples that cannot be assigned to any of the other categories. Each workstation employs a predefined set of growth media, optimised to culture the agents of infection with high sensitivity.

The hospital hygiene department specifically screens for multidrug-resistant pathogens in order to take actions which prevent nosocomial transmission of these. These samples are cultured primarily on selective media containing antibiotics, enabling the growth of resistant strains only.

Growth media have an impact on the bacteria’s proteome and thereby on the MALDI-TOF MS spectrum^57^. In order to avoid that our classifiers recognise media specific characteristics in the MALDI-TOF mass spectra from the selective media instead of media independent signatures of non-susceptible bacterial strains, we excluded samples that were collected for the hospital hygiene department from *DRIAMS-A* for further analysis. The individual sample sizes per workstation and their predictive performance from MALDI-TOF mass spectra is given in **Suppl. Tab. 5**.

### Patient case identification

For *DRIAMS-A*, a *clinical cas*e was defined as a unique hospital stay, i.e. the timeframe between the hospital entry and exit of a patient. If a patient was treated at the hospital in 2015 and again in 2018, these were defined as two separate cases. For the retrospective clinical analysis, infections with different bacterial species and different patient isolation materials during the same hospital stay were regarded as different entities, as different species might require different antibiotic therapies.

For *DRIAMS-B*, *DRIAMS-C* and *DRIAMS-D*, no information regarding *clinical cases* was provided and therefore not considered during analysis.

### Antimicrobial nomenclature

Since the naming scheme of antimicrobial drugs was not consistent between sites, due to different spelling variants or the use of several names for the same drug (e.g. cotrimoxazol and trimethoprim/sulfamethoxazole describe the same drug), we unified them during preprocessing. Specifically, we anglicised the names of antimicrobial drugs from their original German version in the DRIAMS ID files as a preprocessing step for our machine learning analysis. Antimicrobial names were specified by adding suffixes to their names. In the following, we explain our suffix nomenclature for DRIAMS-A: (i) ‘high level’: According to EUCAST, MIC breakpoints vary with the dosage of Gentamycin (standard: 5mg / kg and high level: 7 mg / kg for intravenous administration); (ii) ‘meningitis’, ‘pneumoniae’, ‘endocarditis’, ‘uncomplicated urinary tract infection (UTI)’: EUCAST includes infection specific breakpoints for these infections (see suffixes for amoxicillin-clavulanic acid, penicillin and meropenem); (iii) ‘screen’: cefoxitin is used to screen for MRSA in clinical routine diagnostic at University Hospital Basel, using the respectively-defined breakpoints by EUCAST; (iv) ‘GRD’: GRD (glycopeptide resistance detection) is a MIC Strip Test used at University Hospital Basel in very rare cases to detect glycopeptide intermediate *S. aureus*; (v) ‘1mg_l’ indicates the concentration of rifampicin, when MIC are measured in liquid culture as it is routinely done for *Mycobacterium tuberculosis* (MTB). These entries of non-MTB species were entered incorrectly into the laboratory information system.

### Dataset characteristics

All medical institutions are located in Switzerland. Microbial samples in the University Hospital Basel-Stadt database (i.e. *DRIAMS-A*) mostly originate from patients located in the city of Basel and its surroundings. Such patients visit the hospital for either out- or inpatient treatment. Samples in the Canton Hospital Basel-Land dataset (i.e. *DRIAMS-B*) primarily originate from the town surrounding the City of Basel. Patients from the Swiss Canton Aargau seek medical care at the Canton Hospital Aarau (*DRIAMS-C*). Viollier (*DRIAMS-D*) is a service provider that performs species identification for microbial samples collected in medical practices and hospitals. Samples originate from private practices and hospitals all over Switzerland.

*DRIAMS-A* to *-D* are datasets that contain data collected in the daily clinical routine. All mass spectra measured in a certain time frame are included. The time frame during which each dataset was collected are as follows:

*DRIAMS-A*: 34 months (11/2015–08/2018)

*DRIAMS-B*: 6 months (01/2018–06/2018)

*DRIAMS-C*: 8 months (01/2018–08/2018)

*DRIAMS-D*: 6 months (01/2018–06/2018)

### Spectral representation

In the *DRIAMS* dataset, we include mass spectra in their raw version without any preprocessing, and binned with several bin sizes. After an initial review of results, a bin size of 3 Da was used for all machine learning analyses in this study. This bin size is small enough to allow for separation of mass peaks (for which the exact mass-to-charge position can vary slightly due to measurement noise), while large enough not to impede computational tractability. The spectra are extracted from the Bruker Flex machine in the Bruker Flex data format. The following preprocessing steps are performed using the R package `MaldiQuant`^58^ version 1.19: (1) the measured intensity is transformed with a square-root method to stabilize the variance, (2) smoothing using the Savitzky-Golay algorithm with half-window-size 10 is applied, (3) an estimate of the baseline is removed in 20 iterations of the SNIP algorithm, (4) the intensity is calibrated using the total-ion-current (TIC), and (5) the spectra are trimmed to values in a 2,000 to 20,000 Da range. For exact parameter values, please refer to the code.

After preprocessing, each spectrum is represented by a set of measurements, each of them described by its corresponding mass-to-charge ratio and intensity. However, this representation results in each sample having potentially a different dimensionality (i.e. cardinality) and different measurements being generally irregularly-spaced. Since the machine learning methods used in this manuscript require their input to be a feature vector of fixed dimensionality, intensity measurements are binned using the bin size of 3 Da. To perform the binning, we partition the m/z axis in the range of 2,000 to 20,000 Da into disjoint, equal-sized bins and sum the intensity of all measurements in the sample (i.e. a spectrum) falling into the same bin. Thus, each sample is represented by a vector of fixed dimensionality, i.e. a vector containing 6,000 features, which is the number of bins the m/z axis is partitioned into. We use this feature vector representation for all downstream machine learning tasks.

### Antimicrobial resistance phenotype binarization

For the machine learning analysis, the values of antimicrobial resistance profiles were binarised during data input to have a binary classification scenario. The categories are based on EUCAST and CLSI recommendations. For tests that report RSI values, resistant (R) and intermediate (I) samples were labelled as class 1, while susceptible (S) samples were labelled as class 0. We grouped samples in the intermediate class together with resistant samples, as both types of samples prevent the application of the antibiotic. In EUCAST v6-v8, the intermediate category shows higher MIC values but due to safety reasons in clinical practice this was usually counted to resistance in order to have a safety buffer in reaching sufficiently high antibiotic drug concentrations.

### Machine learning methods

For AMR classification, we used a set of state-of-the-art classification algorithms with different capabilities. It included (1) logistic regression, (2) LightGBM^59^, a modern variant of gradient-boosted decision trees, and (3) a multi-layer perceptron deep neural network (MLP). For LightGBM, we use the official implementation in the `lightgbm` package, while we use the scikit-learn package for all other models^60^. These models cover a large spectrum of modern machine learning techniques, with logistic regression representing a well-understood algorithm from statistical learning theory, whose training process can be regularised. LightGBM, by contrast, represents a modern variant of tree-based (i.e. *ensemble*) learning algorithms, focussing specifically on good scalability properties while maintaining high accuracy. Finally, MLPs constitute a simple example of deep learning algorithms. While they have the highest complexity in terms of compute resources and data requirements than the aforementioned models, deep learning methods can be effective in uncovering complex relationships between input variables, and their capability to be trained end-to-end can prove beneficial in other tasks beyond mere classification.

For each antibiotic, all samples with a missing AMR profile were removed and the machine learning pipeline was applied to the reduced dataset. Samples were randomly split into a training dataset comprising 80% of the samples and a test dataset with the remaining 20%, while stratifying for (1) the class, (2) the species, and (3) the patient case number of the samples. The latter ensures that a sample with a specific clinical case is either part of the train dataset, or the test dataset, but not both. This step ensures that sample measurements of the same infection (that are likely very similar to each other) are causing information leakage from train to test. This is slightly unusual in standard machine learning setups, which typically only require stratification by a single class label, but crucial for our scenarios to guarantee similar prevalence values. To select an appropriate model configuration for a specific task, we employ 5-fold cross-validation; in case an insufficient number of samples is available, our implementation falls back to a 3-fold cross-validation on the training dataset to optimise the respective hyperparameters. The hyperparameters are model-specific (see below for more details), but always include the choice of an optional standardisation step (in which feature vectors are transformed to have zero mean and unit variance). To determine the best-performing hyperparameter set, we optimised the area under the ROC curve (AUROC). This metric is advantageous in our scenario, as it is not influenced by the class ratio and summarises the performance of correct and incorrect susceptibility predictions over varying classification score thresholds. Having selected the best hyperparameters, we retrain each model on the full training dataset, and use the resulting classifier for all subsequent predictions. Our hyperparameter grid is extensive, comprising, for example, the choice of different logistic regression penalties (L1, L2, no penalty), the choice of scaling method (standardisation or none), and regularisation parameters (*C* ∈ {10^-3^, 10^-2^, ··· 10^2^, 10^3^}). For more details, please refer to our code (`models.py`).

We implemented all models in Python and published them in a single package, which we modelled after scikit-learn, a powerful library for machine learning in Python.

### Evaluation metrics

We report AUROC as the main metric of performance evaluation. The datasets of most antibiotics under consideration exhibit a high class imbalance (20 out of 42 antibiotics show a resistant/intermediate class ratio <20% or >80%). AUROC is invariant to the class ratio of the dataset and therefore permits a certain level of comparability between antibiotics with different class ratios. A pitfall of reporting AUROC in the case of unbalanced datasets, however, is that it does not reflect the performance with respect to precision (or positive predictive value). Therefore, the AUROC can be high while precision is low, and it is tempting to be overly optimistic about the AUROC values. To account for this bias, we additionally report the area under the precision-recall curve (AUPRC); this metric is not used during the training process, though.

Two other metrics commonly used in clinical research are sensitivity and specificity. Sensitivity measures to what extent positives are recognized and specificity measures how well negatives are detected as such. Analogous to the ROC curves, we show sensitivity vs. specificity curves to illustrate the tradeoff between both metrics. Please note commonalities to other metrics: *sensitivity*, *recall* and *true positive rate* are synonyms and all correspond to the same metric; specificity is a counterpart to the false positive rate, we have *true positive rate* = 1 − *specificity*.

#### Connection to confusion matrix

All of the metrics we employ here can be derived from the counts within a confusion matrix. The number of true positives (TP) is the number of test data points that are classified as resistant or intermediate by the classifier and confirmed resistant or intermediate in a phenotypic antimicrobial susceptibility test. True negatives (TN) is the number of test data points correctly classified as belonging to the susceptibility class. The false positive count (FP) refers to the number of susceptible test data points classified to be resistant/intermediate, whereas false negatives (FN) are the number of resistant/intermediate test data points classified to be susceptible. The area under the receiver operating characteristic (AUROC) shows the true positive rate (TP/TP+FN) against the false positive rate (FP/FP+TN). The AUPRC, as well as the AUROC, is traditionally reported on the minority class. In our scenario, however, while the minority class is the resistant class in most cases, this is not consistent and for some antibiotics more samples of the resistant will be present. The precision-recall curve shows the recall (TP/TP+FN) against the precision (TP/TP+FP). A perfect classifier would show a performance of 1.0 for each AUROC and AUPRC. The performance of a random classifier would be 0.5 for AUROC and percentage of samples of the positive (susceptible) class for AUPRC. As mentioned above, sensitivity and susceptibility are synonyms for recall and precision, respectively.

### Shapley values for interpretability analysis

Next to the overall predictive performance of an algorithm, a crucial aspect of statistical learning is the understanding of *why* a certain class was predicted for a given sample. This necessitates more information about a classifier, such as confidence scores or feature importance values. The Shapley values provide a ‘one-size-fits-all’ solution to this problem: following concepts from game theory, each feature used to obtain a specific prediction can be assigned its overall contribution to said prediction (or ‘payout’ in the parlance of game theory). This makes it possible to assess to what extent certain bins—peaks in the mass spectrum—influence the prediction. Moreover, the Shapley values are capable of assessing *directed* contributions, i.e. the contribution of a low or high value in a specific feature can be incorporated into the calculation of its overall contribution. Thus, Shapley values represent one cornerstone of the growing field of ‘explainable artificial intelligence’, which endeavours to include the human user in the decision-making loop of a machine learning algorithm.

In order to improve the interpretability of our classifiers, we calculated Shapley values using the `shap` package. This package directly supports the explanation of many common machine learning techniques. We used the standard algorithms of the `shap` package to explain the outputs of our logistic regression and LightGBM models. For the MLP, the use of gradient-based explanation techniques turned out to be impossible because of the large memory requirements of the algorithms. We therefore opted to follow common practice and subsample the input data set, reducing it to 50 barycentres, i.e. samples that express most of the variability in the data, via k-means clustering. This enabled us to obtain per-sample Shapley values that contain the relevance of individual features with respect to the overall output of the model.

## Acknowledgements

We thank Olivia Grüninger, Josiane Reist, Yoann Mahiddine, Daniela Lang, Clarisse Straub, and Magdalena Schneider for excellent technical assistance in accessing the mass spectra and antimicrobial resistance data at the University Hospital Basel. We further thank Belén Rodríguez-Sánchez (Hospital General Universitario Gregorio Marañón, Madrid, Spain), Helena Seth-Smith (USB), Carole Kaufmann (USB), and Valentin Pflüger (MabritecAG, Riehen, Switzerland) for valuable advice, technical consulting, and feedback throughout the project. We thank Dexiong Chen (ETHZ) and Jacques Schrenzel (HUG) for proofreading the manuscript and providing valuable feedback.

## Ethical approval

This study received ethical approval through the local ethical review board (EKNZ number: IEC 2019-00729, 2019-01860 and 2019-00748).

## Author Contributions

C.W., B.R., F.L.-L., K.B. designed machine learning experiments; C.W., B.R. implemented all experiments of the machine learning analysis; A.E., A.C., K.K.S. organised data collection; A.C., S.G., O.U., C.L., M.O. extracted clinical data; A.C, C.W. implemented pre-processing of datasets *DRIAMS-A*, *DRIAMS-B*; C.W. implemented pre-processing of datasets *DRIAMS-C* and *DRIAMS-D*; M.O., A.E. provided feedback on clinical implications of resistance predictions; C.W., B.R., A.C. designed all display items; C.W., B.R., A.C., K.B., A.E. wrote the manuscript with the assistance and feedback of all the other co-authors. K.B., A.E. conceived and supervised the study.

## SUPPLEMENT

**Supplemental Figure 1:**
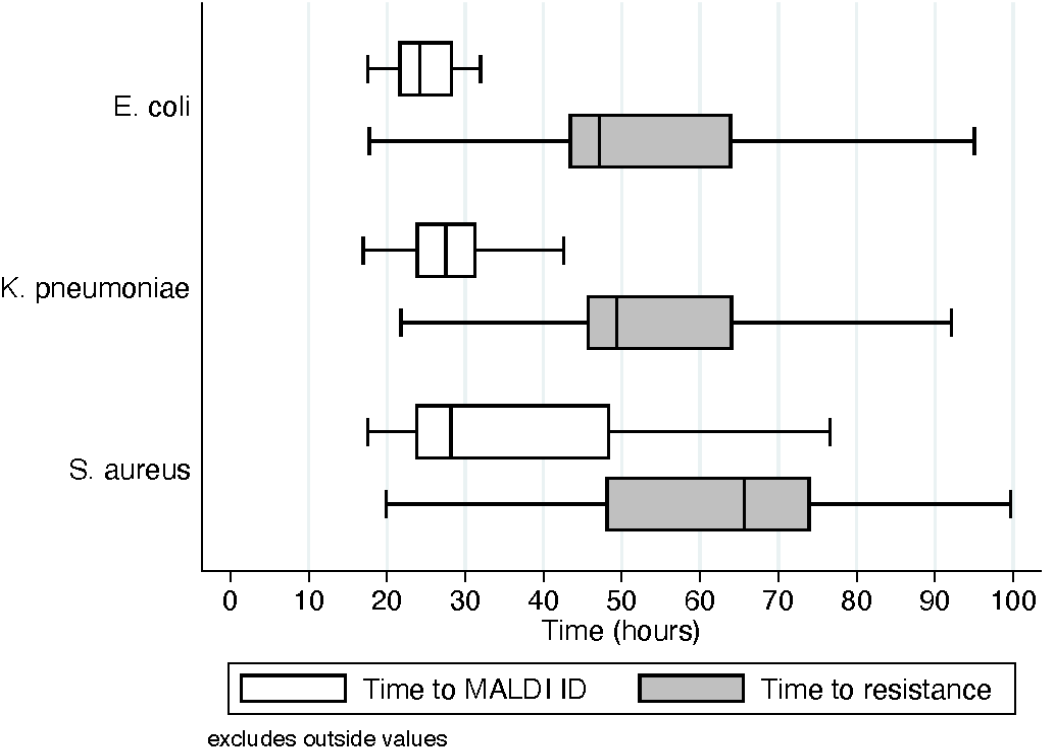
Time from the entry of a patient sample at the diagnostic laboratory at the *DRIAMS-A* collection site to species identification by MALDI-TOF MS and phenotypic resistance testing for three clinical relevant species: *E. coli* (n=54)*, K. pneumoniae* (n=66), and *S. aureus* (n=57). Boxplot shows median and interquartile time ranges in hours, whiskers indicate adjacent values^61^.

**Supplemental Figure 2:**
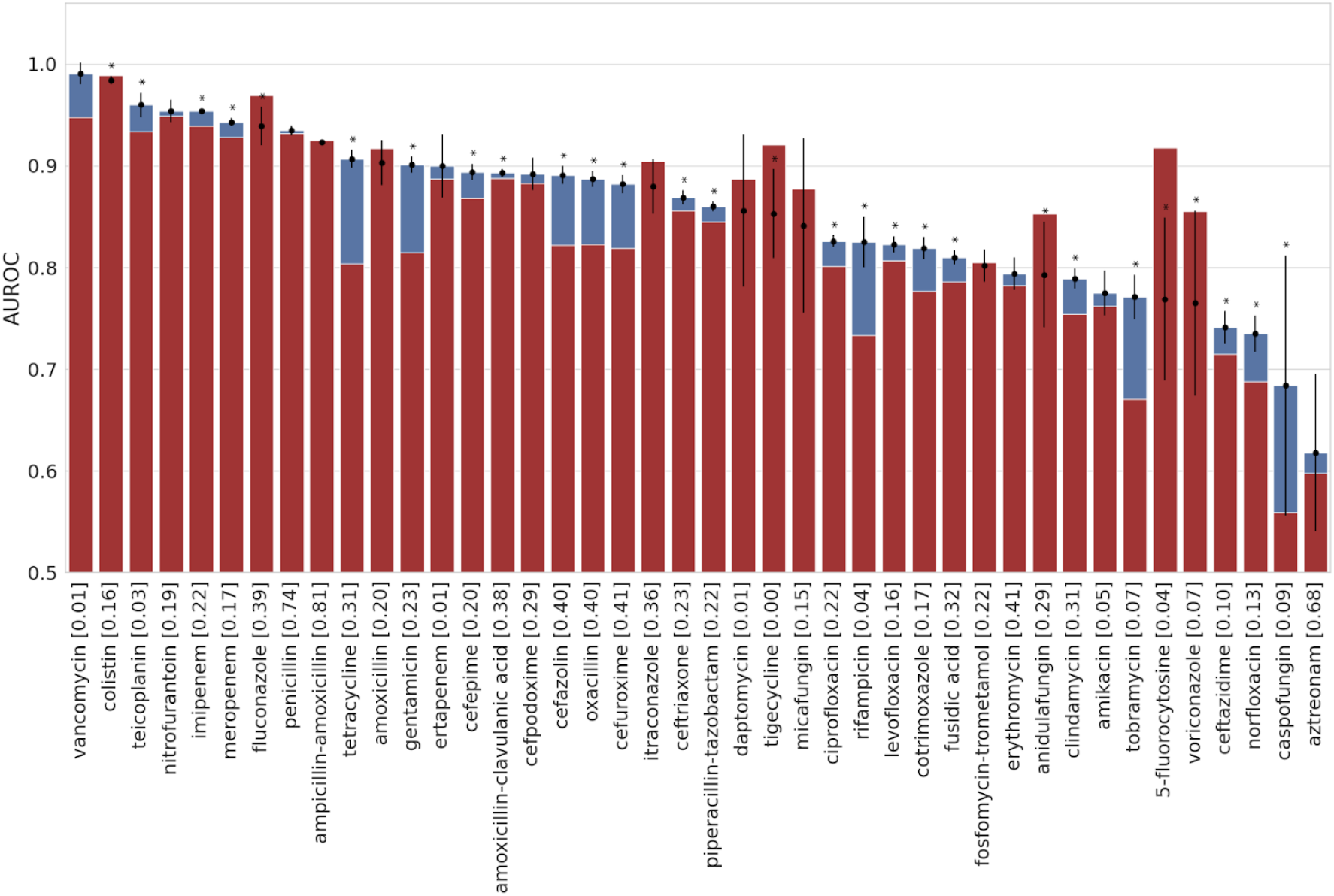
Improved antimicrobial resistance prediction based on MALDI-TOF mass spectra combining all species compared to species information alone. AUROC values of logistic regression classifiers trained on data combining all samples with labels available for each antimicrobial prediction task in *DRIAMS-A.* The blue bars depict predictive performance using spectral data as features. The red bars show the predictive performance when using species label information only. The fractions of resistant/intermediate samples in the training data are indicated in brackets after the antibiotic name. Reported metrics and error bars are the mean and standard deviation of 10 repetitions with different random train-test-splits.

**Supplemental Figure 3:**
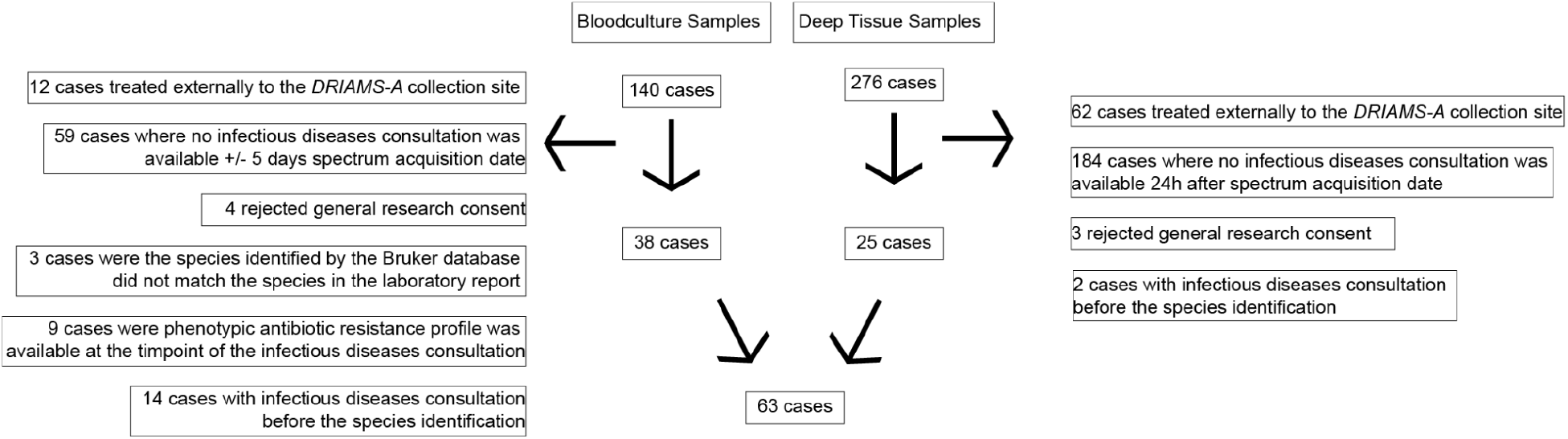
Flowchart inclusion of cases into the retrospective clinical study. We reviewed 416 clinical cases which had a severe bacterial infection with *K. pneumoniae*, *E. coli* or *S. aureus* between April and August 2018. Cases were excluded if (i) cases were treated external to the *DRIAMS-A* collection site, (ii) no consultation note by a infectious diseases specialist was available within 5 days (for cases with a positive blood culture) or 1 day (for cases with a positive deep tissue sample), (iii) the general research consent was rejected, (iv) the species identified by the Bruker database did not match the species in the laboratory report, (v) the antibiotic resistance profile was already present at the at the time of the infectious diseases consultation and (vi) if the consultation note was written without the knowledge of the species identity. 63 clinical cases were included.

**Supplemental Figure 4:**
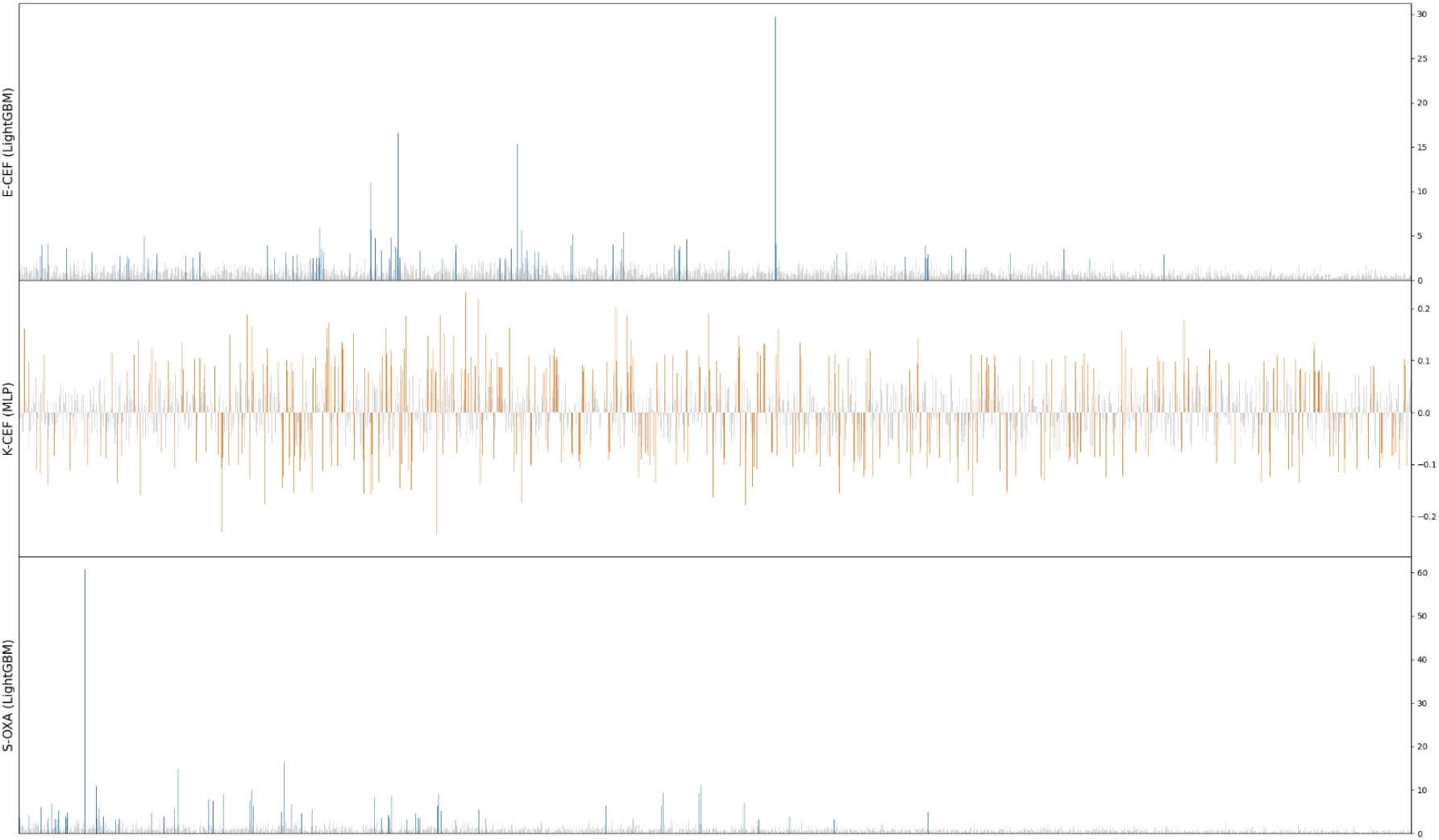
Barplot of feature importances of LightGBM and MLP model. Importance values larger than two times absolute standard deviation are colored in either blue (LightGBM) or orange (MLP). The sign of each feature importance value of the MLP model indicates the association with the positive (positive sign) or negative class. The LightGBM values indicate the contribution to the prediction without direction of association. All three models indicate that a large number of features are relevant for an accurate antimicrobial resistance prediction. The scenario abbreviations follow **Fig. 2A**.

**Supplemental Figure 5:**
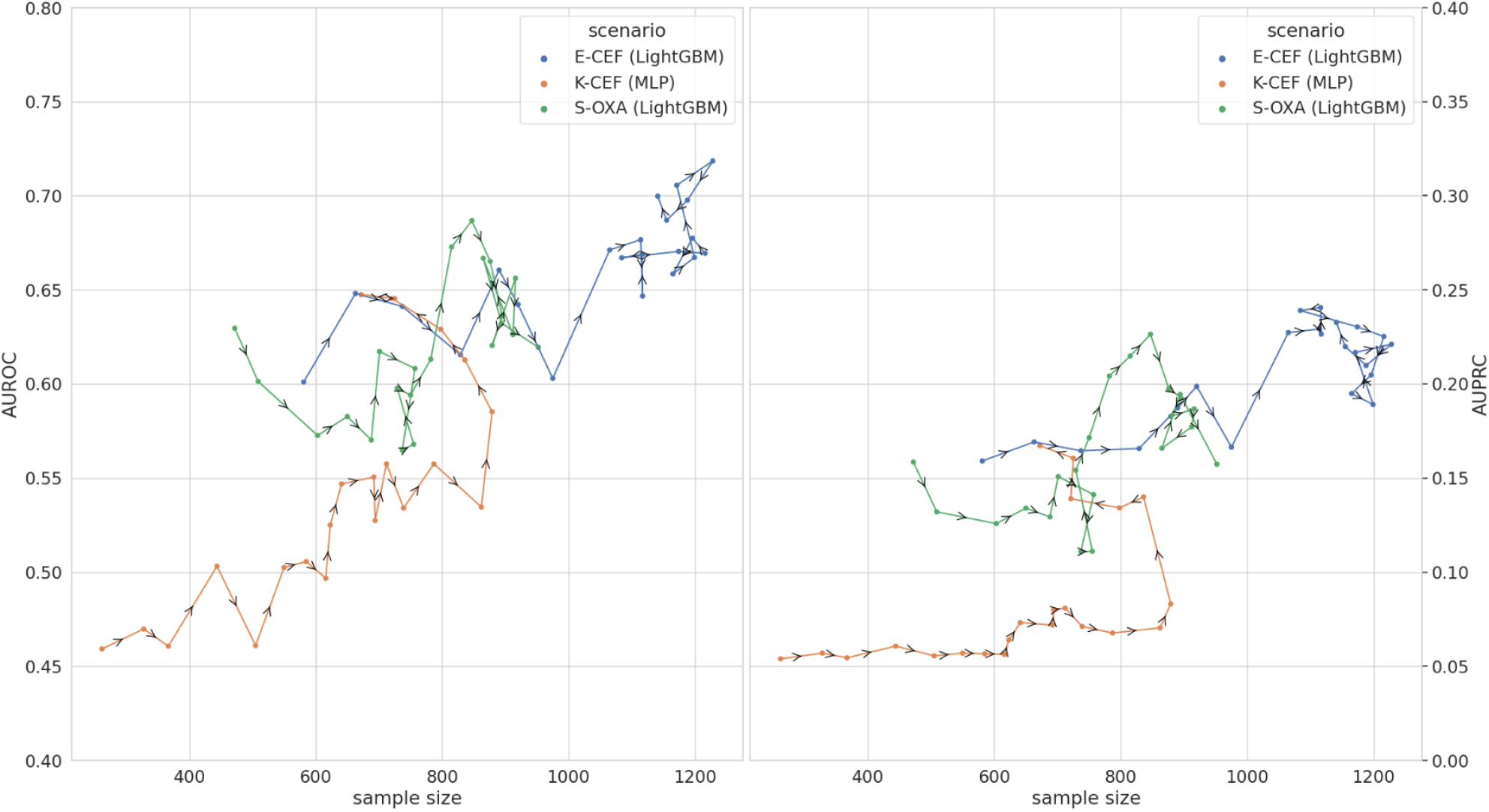
Temporal validation including sample size of training window. The timepoints correspond to points in **Fig. 4** and arrow directions indicate time progression. With time progression both the trends in sample size per 8-month time window and the predictive performance increase. The scenario abbreviations follow **Fig. 2A**.

**Supplemental Table 1:**
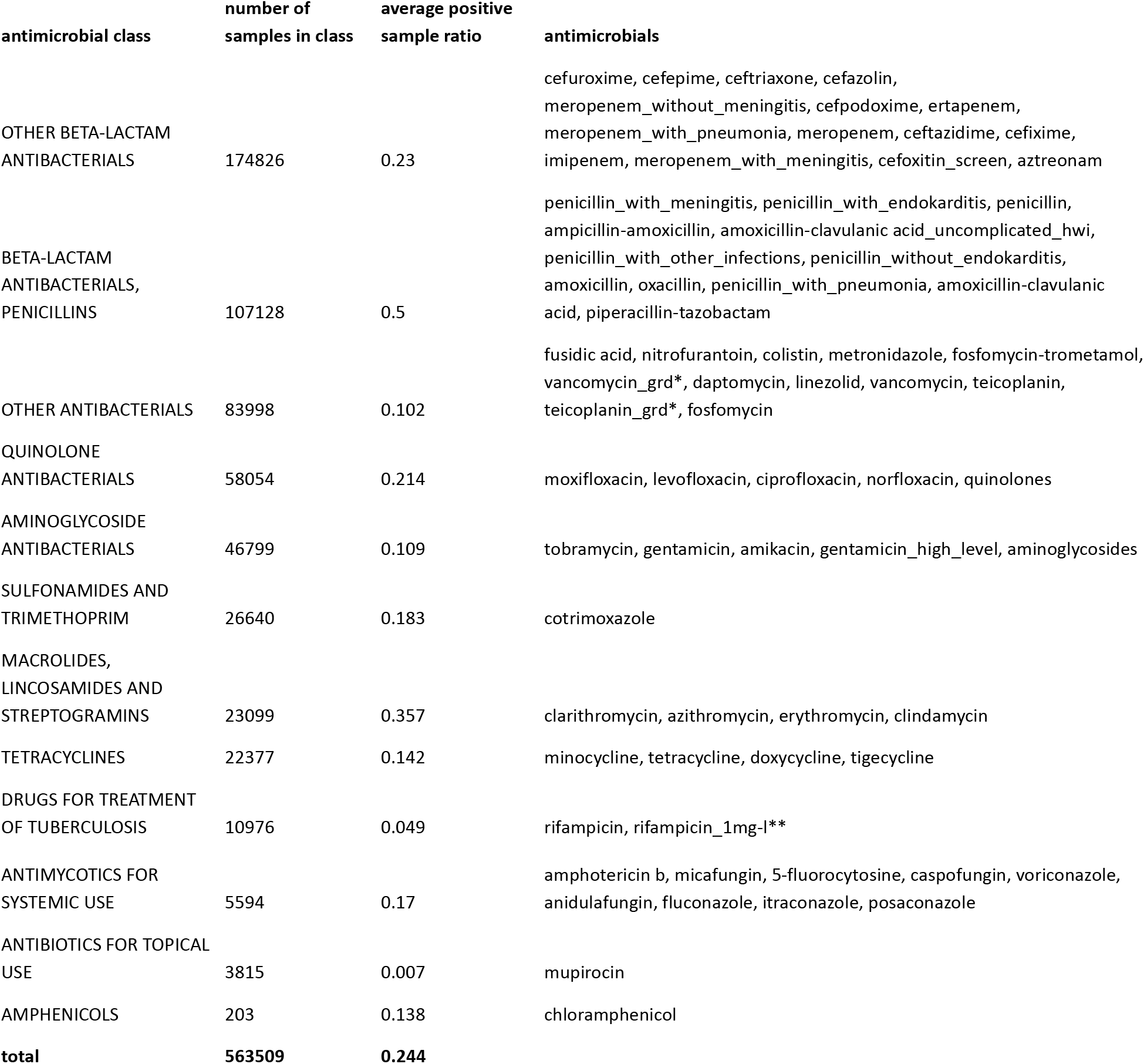
Categorisation into antimicrobial class, average number of spectra and average resistance positive class ratio of 71 antimicrobials contained in *DRIAMS-A*. The values for each individual antimicrobial can be found in **Suppl. Tab. 2**. * vancomycin_grd and teicoplanin_grd is a MIC Strip Test used at University Hospital Basel in very rare cases to detect glycopeptide intermediate *S. aureus*; ** ‘1mg_l’ indicates the concentration of rifampicin, when MIC are measured in liquid culture as it is routinely done for *Mycobacterium tuberculosis* (MTB). These entries of non-MTB species were entered incorrectly into the laboratory information system.

**Supplemental Table 2:**
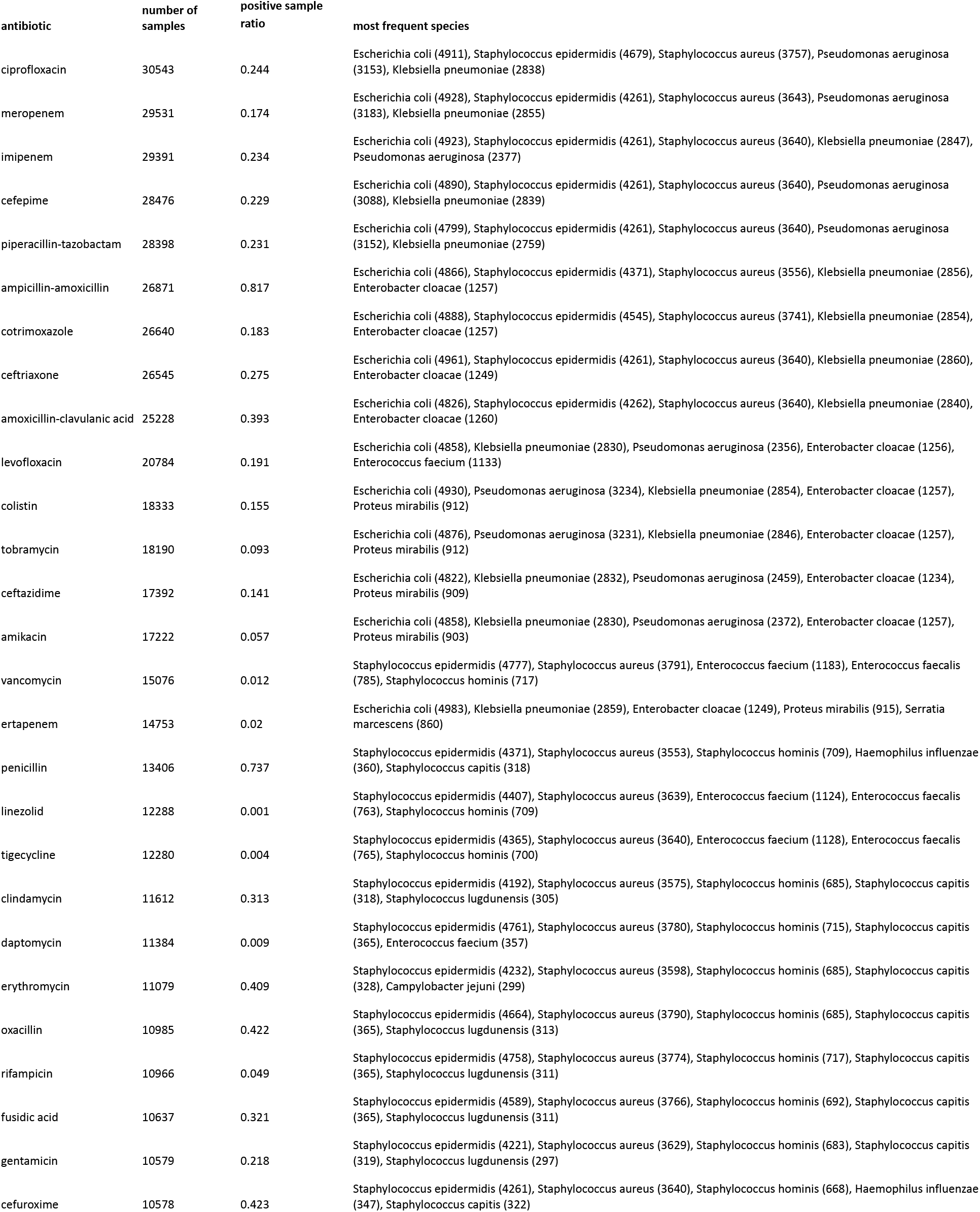
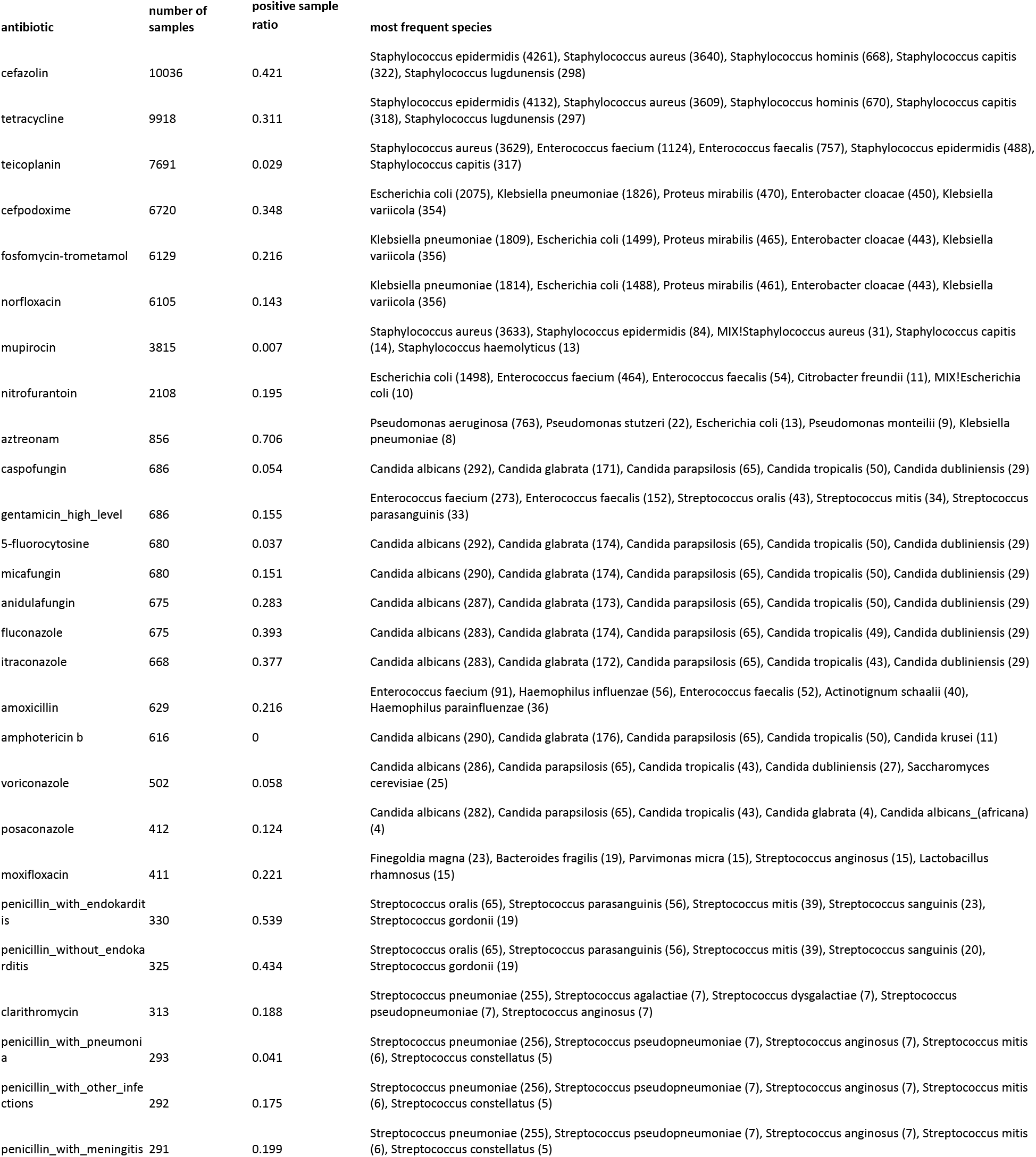
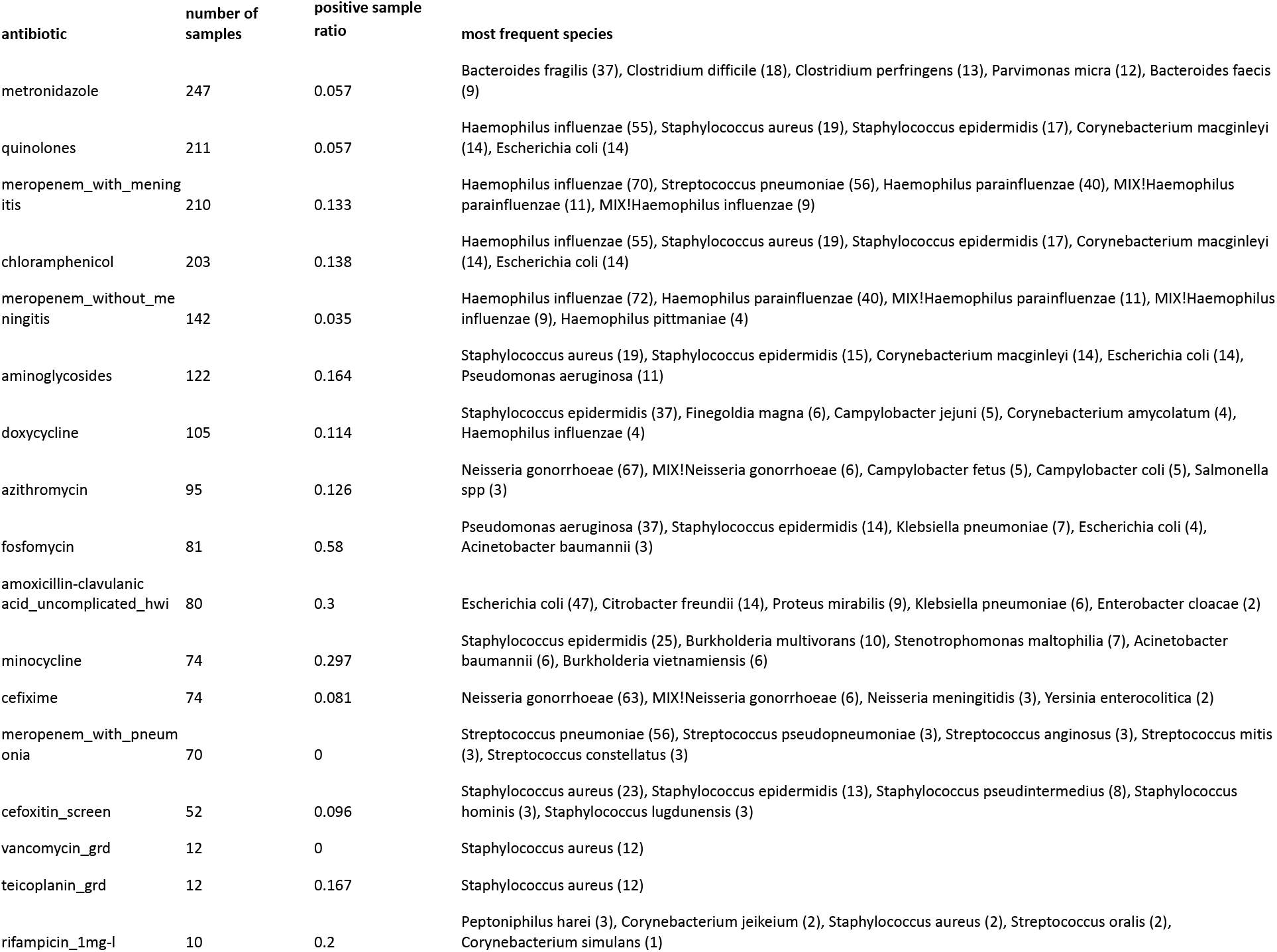
Complete list of antibiotics contained in *DRIAMS-A*, including (i) the number of MALDI-TOF mass spectra for which the antimicrobial resistance was determined, (ii) the resistance class ratio among these samples (positive sample ratio) and (iii) the five most frequent species among these samples with the number of samples indicated in brackets. * Antibiotic resistance category is inferred from oxacillin resistance. ** Antibiotic resistance category is inferred from ciprofloxacin resistance. *** Antibiotic resistance category for this species is automatically set to the resistant category. **** Part of a species complex. ***** Breakpoints different according to clinical presentation (see EUCAST guidelines). + vancomycin_grd and teicoplanin_grd is a MIC Strip Test used at University Hospital Basel in very rare cases to detect glycopeptide intermediate *S. aureus*; ++ ‘1mg_l’ indicates the concentration of rifampicin, when MIC are measured in liquid culture as it is routinely done for *Mycobacterium tuberculosis* (MTB). These entries of non-MTB species were entered incorrectly into the laboratory information system.

**Supplemental Table 3:** Union experiment results for *DRIAMS-C* and *DRIAMS-D* in AUPRC. **A.** Results for test site *DRIAMS-C*. The predictive performance benefits from combining the target site train data from *DRIAMS-C* with the large *DRIAMS-A* datasets, compared to training one either train site alone. For two scenarios adding *DRIAMS-B* further improves the performance slightly. **B.** Results for test site *DRIAMS-D*. The behaviour differs from the other test sites in that training exclusively on the test site DRIAMS-D leads to the best predictive performance. No samples for *S. aureus* (oxacillin) available for *DRIAMS-D*.

| **A. Test site: DRIAMS-C (class 1 ratio 16.3%)** |  |  | **Train site:**<br>**DRIAMS-A** | **Train site:**<br>**DRIAMS-C** | **Train sites:**<br>**DRIAMS-A**<br>**DRIAMS-C** | **Train sites:**<br>**DRIAMS-A**<br>**DRIAMS-B**<br>**DRIAMS-D\*** | **Train sites:**<br>**DRIAMS-A**<br>**DRIAMS-B**<br>**DRIAMS-C**<br>**DRIAMS-D\*** |
| --- | --- | --- | --- | --- | --- | --- | --- |
| **species** | **antibiotic** | **model** |  |  |  |  |  |
| *E. coli* | ceftriaxone | LightGBM | 0.31 ± 0.06 | 0.34 ± 0.06 | 0.39 ± 0.05 | 0.35 ± 0.06 | **0.40 ± 0.06** |
| *K. pneumoniae* | ceftriaxone | MLP | 0.34 ± 0.09 | 0.42 ± 0.11 | 0.42 ± 0.11 | 0.25 ± 0.07 | **0.44 ± 0.09** |
| *S. aureus* | oxacillin | LightGBM | 0.21 ± 0.09 | 0.08 ± 0.02 | **0.27 ± 0.12** | 0.21 ± 0.12 | 0.27 ± 0.14 |
| **B. Test site: DRIAMS-D (class 1 ratio 10.0%)** |  |  | **Train site:**<br>**DRIAMS-A** | **Train site:**<br>**DRIAMS-D** | **Train sites:**<br>**DRIAMS-A**<br>**DRIAMS-D** | **Train sites:**<br>**DRIAMS-A**<br>**DRIAMS-B**<br>**DRIAMS-C** | **Train sites:**<br>**DRIAMS-A**<br>**DRIAMS-B**<br>**DRIAMS-C**<br>**DRIAMS-D** |
| **species** | **antibiotic** | **model** |  |  |  |  |  |
| *E. coli* | ceftriaxone | LightGBM | 0.29 ± 0.09 | **0.48 ± 0.05** | 0.43 ± 0.06 | 0.34 ± 0.08 | 0.43 ± 0.07 |
| *K. pneumoniae* | ceftriaxone | MLP | 0.09 ± 0.03 | **0.23 ± 0.06** | 0.21 ± 0.05 | 0.11 ± 0.04 | 0.20 ± 0.04 |

**Supplemental Table 4:** Highly ranked feature bins which have been identified in previous studies. We performed a thorough literature research including studies which reported peaks falling into feature bins receiving high weight of our classifier and report relevant peaks. ‘NA’: not available as this study did not determine new discriminatory peaks, but focussed on identifying previously known peaks^44^. ‘Target’ which subgroups within the species were aimed to be discriminated against.

| species | feature bin [m/z] | rank (feature importance) | rank (Shapley values) | Ref: exact m/z value given | Ref: method discriminatory peak identification | Ref: number of strains | Ref: target |
| --- | --- | --- | --- | --- | --- | --- | --- |
| *S. aureus* | 2759-2762 | 14 | 12 | 2762<sup>50</sup> | *Genetic algorithm via ClinProTools* | 290 | Dominant lineages within MRSA (ST5) |
|  |  |  |  | 2760<sup>48</sup> | *Support Vector Machine* | 160 | Discrimination between MSSA and MRSA |
| *S. aureus* | 3005-3008 | 13 | 7 | 3007<sup>44</sup> | NA | 401 | Main Clonal Lineages (CC1) |
| *S. aureus* | 3890-3893 | 4 | 3 | 3891<sup>45</sup> | Visual Examination | 600 | CC5, CC97 |
|  |  |  |  | 3891<sup>44</sup> | NA | 401 | Main Clonal Lineages (CC5, CC25) |
| *S. aureus* | 4508-4511 | 19 | 14 | 4511<sup>49</sup> | Visual examination | 85 | Major lineages within MRSA (CC45, CC30) |
|  |  |  |  | 4511<sup>44</sup> | NA | 401 | CC30, CC45, CC398, ST88 |
| *S. aureus* | 4514-4517 | 12 | 15 | 4514<sup>42</sup> | *Supervised neural network via ClinProTools* | 82 | MRSA Clonal Complexes (CC398) |
| *S. aureus* | 4640-4643 | 2 | 2 | 4641<sup>49</sup> | Visual examination | 85 | Major lineages within MRSA (CC8, CC22) |
|  |  |  |  | 4641<sup>48</sup> | *Support Vector Machine* | 160 | Discrimination between MSSA and MRSA |
| *S. aureus* | 5003-5006 | 3 | 4 | 5002<sup>49</sup> | Visual examination | 85 | Major lineages within MRSA (CC22) |
|  |  |  |  | 5004<sup>42</sup> | *Supervised neural network via ClinProTools* | 82 | MRSA Clonal Complexes (CC22) |
|  |  |  |  | 5002<sup>45</sup> | Visual examination | 600 | CC22 |
|  |  |  |  | 5002<sup>44</sup> | NA | 401 | Main Clonal Lineages (CC22) |
| *S. aureus* | 5432-5435<br>5435-5438 | 6, 7 | 6, 5 | 5437<sup>49</sup> | Visual examination | 85 | Major lineages within MRSA (CC5, CC45, CC22, CC8, ST1, ST15, ST80) |
|  |  |  |  | 5440<sup>47</sup> | Support Vector Machine | 626 | MSSA CC98 |
| *E. coli* | 8501-8504 | 10 | 8 | 8496<sup>62</sup> | Support Vector Machine | 109 | ST131 |
| *E. coli* | 8444-8447<br>8447-8450<br>8450-8453 | 9, 5, 3 | 7, 5, 2 | 8448<sup>54</sup> | *Peak Statistic Calculation* in ClinProTools | 197 | ST131 |
| *E. coli* | 11780-11783 | 1 | 1 | 11783<sup>54</sup> | *Peak Statistic Calculation* in ClinProTools | 197 | ST131 |

**Supplemental Table 5:** Overview of sample distribution and distinguishability over the workstations in *DRIAMS-A.* For each scenario, the number of samples and the percentage of the total number of samples is stated for each workstation. Additionally, a multi-class logistic regression classifier was trained to predict the workstation based on the MALDI-TOF spectrum, and the predictive performance is reported by the average recall. Generally, we observe a high recall for all workstations if a substantial sample size is present; particularly for hospital hygiene and blood, substantiating prior knowledge that the growth medium can be detected from the MALDI-TOF mass spectra (see **Methods**).

| species | antibiotic |  | hospital<br>hygiene | blood | deep tissue | genital | respiratory | stool | urine | varia | total |
| --- | --- | --- | --- | --- | --- | --- | --- | --- | --- | --- | --- |
| *E. coli* | ceftriaxone | n | 659 | 1190 | 1073 | 24 | 364 | 5 | 1473 | 173 | 4961 |
|  |  | % | 13.3 | 24.0 | 21.6 | 0.5 | 7.3 | 0.1 | 29.7 | 3.5 | 100.0 |
|  |  | recall | 88.8 ± 0.3 | 77.8 ± 0.7 | 57.6 ± 1.4 | 0.0 ± 0.0 | 0.0 ± 0.0 | 0.0 ± 0.0 | 83.5 ± 0.5 | 0.0 ± 0.0 |  |
| *K. pneumoniae* | ceftriaxone | n | 229 | 273 | 204 | 5 | 268 | 15 | 1790 | 76 | 2860 |
|  |  | % | 8.0 | 9.6 | 7.1 | 0.2 | 9.4 | 0.5 | 62.6 | 2.7 | 100.0 |
|  |  | recall | 79.1 ± 0.5 | 62.4 ± 0.4 | 0.0 ± 0.0 | 0.0 ± 0.0 | 46.5 ± 0.8 | 0.0 ± 0.0 | 99.6 ± 0.1 | 3.8 ± 2.6 |  |
| *S. aureus* | oxacillin | n | 379 | 708 | 1356 | 34 | 517 | 0 | 187 | 609 | 3790 |
|  |  | % | 10.0 | 18.7 | 35.8 | 0.9 | 13.6 | 0.0 | 4.9 | 16.1 | 100.0 |
|  |  | recall | 89.9 ± 0.5 | 52.3 ± 2.2 | 90.8 ± 0.7 | 0.0 ± 0.0 | 33.4 ± 1.0 | NA | 1.8 ± 1.5 | 2.2 ± 2.0 |  |

